# Starvation-induced cell fusion and heterokaryosis frequently escape imperfect allorecognition systems to enable parasexual interactions in an asexual fungal pathogen

**DOI:** 10.1101/2021.05.19.444787

**Authors:** Vasileios Vangalis, Ilya Likhotkin, Michael Knop, Milton A. Typas, Ioannis A. Papaioannou

**Author notes:** Author for correspondence: Ioannis A. Papaioannou.

## Abstract

- Asexual fungi include important pathogens of plants and other organisms, and their effective management requires understanding of their evolutionary dynamics. Genetic recombination is critical for species adaptability and could be achieved via heterokaryosis and the parasexual cycle in asexual fungi. Here, we investigate the extent and mechanisms of heterokaryosis in the asexual plant pathogen *Verticillium dahliae*.
- We used live-cell imaging and genetic complementation assays of tagged *V. dahliae* strains to analyze the extent of nonself vegetative fusion, heterokaryotic cell fate and nuclear behavior. An efficient CRISPR/Cas9-mediated system was developed to investigate the involvement of autophagy in heterokaryosis.
- Under starvation, nonself fusion of germinating spores occurs frequently regardless of the previously assessed vegetative compatibility of the partners. Supposedly “incompatible” fusions often establish viable heterokaryotic cells and mosaic mycelia, where nuclei can engage in fusion or transfer of genetic material. The molecular machinery of autophagy has a protective function against destruction of “incompatible” heterokaryons.
- Our results suggest an autophagy-mediated trade-off between parasexual interactions for genetic exchange and allorecognition systems possibly for mycelial protection from parasitic elements. Our study reveals unexpected capacity for heterokaryosis in *V. dahliae* and suggests, therefore, important roles of parasexuality in the evolution of asexual fungi.

## Introduction

Generation of genetic and phenotypic diversity is of utmost importance for the survival and adaptability of species over evolutionary timescales (Booy *et al*., 2000; Frankham, 2005). Genetic reshuffling in meiosis provides a constant supply of genetic substrates for natural selection and enables efficient purging of deleterious mutations from sexual populations (Fisher, 1930; Muller, 1932). A complete lack of recombination in organisms such as obligate asexual species can result in the accumulation of problematic alleles, a phenomenon known as “Muller’s Ratchet” (Felsenstein, 1974). Some source of recombination is, therefore, usually deemed essential for the long-term survival of species.

In earlier days of fungal genetics, an alternative mechanism for genetic recombination, termed the parasexual cycle, was described in fungi (Pontecorvo & Roper, 1952). This refers to a combination of non-sexual processes that lead to genetic diversification through somatic cell and nuclear fusion, followed by mitotic recombination during ploidy reduction (i.e. haploidization) (Pontecorvo & Roper, 1952; Pontecorvo, 1956). Meiotic sexual cycles may have evolved from ancestral parasexual processes during the early eukaryotic evolution (Goodenough & Heitman, 2014). Variations of the parasexual cycle are present in several and diverse extant eukaryotes, i.e. many filamentous fungi and yeasts (Tinline & MacNeill, 1969; Strom & Bushley, 2016; Anderson *et al*., 2019; Zhao *et al*., 2020), protists (Gaunt *et al*., 2003; Carpenter *et al*., 2012) and mammalian cells, where it can contribute to the phenotypic plasticity of tumor cell populations (Miroshnychenko *et al*., 2021).

The parasexual cycle is normally initiated in heterokaryons, i.e. cells that host genetically distinct nuclei in common cytoplasm. Heterokaryosis in fungi often arises from somatic non-self fusion (or anastomosis) between hyphae or spores of different strains (Roca *et al*., 2005; Mela *et al*., 2020). However, fusion between different individuals and establishment of viable heterokaryons are believed to be largely prevented in nature by non-self recognition systems, a phenomenon described as vegetative (or heterokaryon) incompatibility (Saupe, 2000; Paoletti, 2016; Daskalov *et al*., 2017). Incompatibility barriers can block cell communication to prevent fusion, or they can trigger post-fusion rejection mechanisms that result in the induction of programmed cell death in the incompatible heterokaryotic cell (Paoletti, 2016; Daskalov *et al*., 2017; Gonçalves & Glass, 2020). The versatile allorecognition mechanisms have been analyzed mostly in the sexual fungi *Neurospora crassa*, *Podospora anserina* and *Cryphonectria parasitica*, where they are believed to function as defense mechanisms against the rapid spread of infectious elements and somatic parasitism (Saupe, 2000; Paoletti, 2016). Fungal strains unable to form viable vegetative heterokaryons because of these barriers are referred to as vegetatively incompatible, whereas compatible strains that can fuse and establish stable heterokaryons are classified into the same Vegetative Compatibility Group (VCG) (Leslie, 1993). Despite the large perceived extent of incompatibility in natural populations (Leslie, 1993; Clutterbuck, 1996), many fungi seem competent in parasexual interactions in the laboratory (Tinline & MacNeill, 1969; Strom & Bushley, 2016; Anderson *et al*., 2019), and the possibly heterokaryosis-mediated horizontal gene transfer has gained increasing attention in the postgenomic era as a driver of fungal evolution (Fitzpatrick, 2012; Soanes & Richards, 2014).

The soil-borne asexual ascomycete *Verticillium dahliae* is the causal agent of Verticillium wilt, a disease that affects a wide range of plants and accounts for significant annual economic losses (Pegg & Brady, 2002). This haploid species is considered strictly asexual and propagates predominantly by clonal expansion (Gurung *et al*., 2014). Despite the apparent lack of sexual activity, *V. dahliae* populations exhibit signs of recombination, which, although not frequent, has the potential to create novel clonal lineages (Atallah *et al*., 2010; Milgroom *et al*., 2014). The parasexual cycle has been described in *Verticillium* species and presents interesting features (Hastie, 1964; Puhalla & Mayfield, 1974; Typas & Heale, 1976). These include high instability of transient diploid nuclei, which are formed within heterokaryons following cell fusion, and frequent mitotic recombination during haploidization, often leading to recombined progeny (Hastie, 1964, 1968; Typas & Heale, 1978). Furthermore, overcoming cell wall-related barriers by protoplast fusion or nuclear microinjection resulted in interspecific heterokaryons between *V. dahliae* and *V. albo-atrum* at frequencies similar to intraspecific combinations (Typas, 1983). Our previous finding that heterokaryosis in *V. dahliae* is occasionally possible even between strains of different VCGs (thus being considered incompatible) (Papaioannou & Typas, 2015) raises the question whether parasexuality in asexual fungi may be more important than previously recognized. We recently established fusion between spores via conidial anastomosis tubes (CATs) as a convenient system for addressing this question (Vangalis *et al*., 2021).

In this work, our analyses of the extent and mechanisms of heterokaryosis in the strictly asexual fungus *V. dahliae* challenge the general assumption of absolute vegetative incompatibility barriers in asexual fungi. Frequent heterokaryon formation and transfer of genetic material even between supposedly “incompatible” partners, under non-selective conditions, suggest that parasexuality plays an important role in genomic and phenotypic diversification. Overall, our study paves the way for the elucidation of non-sexual genetic interactions in asexual fungi and the evolutionary significance of these processes in adaptation and survival.

## Materials and Methods

### Strains and culture conditions

All *V. dahliae* strains and plasmids constructed and used in this study are listed in Table S1 and Table S2, respectively. Their background wild-type isolates were previously classified into Vegetative Compatibility Groups (VCGs) based on traditional complementation assays between nitrate non-utilizing (*nit*) mutants (Papaioannou *et al*., 2014; Papaioannou & Typas, 2015; Vangalis *et al*., 2021). Preparation, handling and maintenance of monoconidial strains, as well as culture media and conditions have been previously described (Papaioannou *et al*., 2013b; Vangalis *et al*., 2021).

### Labeling of strains with fluorescent markers

Nuclei of *V. dahliae* strains were labeled with either sGFP- or mCherry-tagged histone H1, according to the methods described in Method S1. To study cytoplasmic mixing following CAT-mediated cell fusion, *V. dahliae* strains were also constructed to express sGFP with cytoplasmic localization (Method S1).

### Analysis of CAT-mediated cell fusion

Our methods for analysis and quantification of CAT-mediated fusion of *V. dahliae* conidia/germlings have recently been described (Vangalis *et al*., 2021). Briefly, fresh conidial suspensions (in water) were always prepared from 7-day-old PDA cultures and diluted in CAT medium (0.75 g l^-1^ β-glycerophosphate disodium salt · 5H_2_O) to a final concentration of 1.0 × 10^6^ conidia ml^-1^. Suspensions were then mixed (1:1), and 100 μl of their mixture were transferred to a well of a 96-well glass-bottom plate (MGB096-1-2-LG-L from Matrical Bioscience, Spokane, WA, USA). The plate was incubated at 24°C (in the dark) for 60 h before imaging. Each pairing was performed in triplicate, and 250-300 fusion events were analyzed per replicate. Staining with 0.005% methylene blue (Sigma-Aldrich, St. Louis, MO, USA) (incubation at 24°C for 5 min) was used for quantifying cell viability of fused cells.

### Microscopy

Image acquisition was performed using a Nikon (Tokyo, Japan) Ti-E epifluorescence microscope equipped with an autofocus system (Perfect Focus System, Nikon), a LED light engine (Spectra X from Lumencor, Beaverton, OR, USA), filter sets 390/18 and 435/48, 469/35 and 525/50, and 542/27 and 600/52 (excitation and emission, respectively; all from Semrock, Rochester, NY, USA, except for 525/50, which was from Chroma Technology, Bellows Falls, VT, USA), and a sCMOS camera (Flash4.0 from Hamamatsu, Honshu, Japan). At least 20 non-overlapping fields of view were imaged per well. Calcofluor white M2R (Sigma-Aldrich, St. Louis, MO, USA) was used at a final concentration of 10 μg ml^-1^ (incubation at 24°C for 5 min) for staining of chitin in cell walls and septae. In time-lapse experiments, the imaging plates were scanned for image acquisition at 10 min-intervals for up to 24 h (exposure time: 20 ms for the blue channel, 50 ms for the green and red channels) and were incubated undisturbed at 24°C (in the dark between image acquisitions). In some experiments we performed *z*-stack (step size = 0.5 μm) time-lapse imaging at each position, for up to 8 h. Image processing was performed using ImageJ (Schindelin *et al*., 2012). Images were adjusted to a uniform contrast across all time points for each experiment, and the maximum intensity projection method was used for processing *z*-stacks.

### Hyphal- and CAT-based complementation assays

We used two complementation methods for assessing the degree of vegetative genetic isolation between *V. dahliae* strains, namely the traditional VCG (hyphal-based) assay and a newly developed assay that permits CAT-mediated fusion of the interacting strains. The complementation assays are described in Method S2.

### Mixed artificial infection of eggplant seedlings

Eggplant seedlings at the one true leaf-stage were drenched with 20 ml of mixed conidial suspensions (5.0 × 10^6^ conidia ml^-1^) or sterile water (mock-infection controls) (Markakis *et al*., 2014). Plants were incubated at 25°C with a 12-h light-dark cycle. Fungal re-isolation was performed from sections along the stem of each treated plant (from eight xylem chips per plant) 40 days after inoculation, according to described procedures (Markakis *et al*., 2014). Three xylem chips from each plant were transferred to acidified PDA as controls, while the remaining five chips were transferred to acidified double-selective PDA (10 μg ml^-1^ hygB and 100 μg ml^-1^ G418) to select for heterokaryotic growth.

### Identification of autophagy genes

The protein sequences of 41 genes involved in autophagy (Table S3) were retrieved from the Saccharomyces Genome Database (https://www.yeastgenome.org) and the Ensembl Fungi database (http://fungi.ensembl.org/index.html) and used as queries in tBLASTn searches of the *V. dahliae* Ls.17 genome (NCBI Genome assembly GCA_000952015.1) to detect homologs of core autophagy genes, applying previously described criteria (Wang *et al*., 2019). For reasons of comparison, we included in Table S3 data for various classes of the fungal phylogeny, adopted from (Wang *et al*., 2019).

### Lithium acetate (LiAc)/heat shock-mediated transformation of *V. dahliae* conidia

At least 10^8^ conidia of each *V. dahliae* strain were harvested from PDA plates, pelleted, resuspended in 30 ml of a phosphate-KCl buffer (0.6 M KCl, 1/15 M KH_2_PO_4_, and 1/15 M Na_2_HPO_4_, pH 5.8; supplemented with 25 mM DTT) and incubated for 2 h at 30°C (50 rpm). Conidia were then washed once with water and once with a 0.1 M LiAc solution, before being resuspended in 0.5 ml of 0.1 M LiAc and incubated at room temperature for 20 min. Aliquots (100 μl) of this suspension were mixed with 5-10 μg of plasmid, 240 μl of 50% PEG-4000 and 35 μl of 1 M LiAc, and they were incubated on ice for 30 min before being transferred to 34°C for 10 min (heat-shock). Each sample was then washed once and plated on PDA medium overlaid with a cellophane sheet. Following incubation at 25°C for 24 h, the sheet was transferred onto PDA supplemented with hygromycin B (15 μg ml^-1^) for selection of transformants.

### CRISPR/Cas9-mediated gene tagging and knockout

The components and the strategy of our CRISPR/Cas9-based method, adopted from (Nødvig *et al*., 2015), are presented in Fig. 5. An AMA1-containing plasmid (pVV27) with the SpCas9 cassette, an *atg8*-specific sgRNA cassette (for targeting *V. dahliae atg8*, VDAG_01225) and the *hph* selection marker was generated by the insertion of two PCR fragments into plasmid pFC332 (Nødvig *et al*., 2015). For this, two PCR fragments were amplified from plasmid pFC334 using the Herculase II Fusion DNA Polymerase with primers containing the protospacer sequence of *atg8* (Table S4). The backbone plasmid pFC332 was linearized with *Pac*I digestion, and the three fragments were assembled into plasmid pVV27 using the NEBuilder HiFi DNA Assembly Master Mix kit. Repair substrates with 1.0 kb-long homologous arms (Fig. 5b,c) were constructed by cloning the *sgfp* or *mCherry* gene (amplified from plasmid pIGPAPA or pMaM330, respectively; Table S2) and the desired amplified flanks from genomic DNA of *V. dahliae* Ls.17 into plasmids pVV25 and pVV26, respectively (Table S2), using the NEBuilder HiFi DNA Assembly Master Mix kit. Repair substrates with 0.1 kb-long homologous arms (Fig. 5b,c) were obtained directly by PCR amplification from plasmid pIGPAPA, pMaM330 or pSD1 (for the *sgfp*, *mCherry* or *neo*^R^ gene, respectively; Table S2) using primers containing the desired homologous arms (Table S4). Co-transformation of *V. dahliae* strains with plasmid pVV27 and a repair substrate was performed using a protoplast-mediated method (Vangalis *et al*., 2020) or the LiAc/heat shock-mediated method. Transformants were screened by PCR and the fluorescently tagged strains were confirmed by microscopy. To induce autophagy in these experiments, we used rapamycin (Sigma-Aldrich, St. Louis, MO, USA) at a final concentration of 500 nM.

### Gene deletion using *Agrobacterium tumefaciens*-mediated transformation (ATMT) and complementation of knockout strains

The *atg1* homolog of *V. dahliae* (VDAG_05745) was deleted by the ATMT-based method (Fig. S3a) that we have previously described (Vangalis *et al*., 2021). Knockout strains were complemented by re-introducing the corresponding wild-type genes (Vangalis *et al*., 2021).

## Results

### Conidial fusion occurs frequently between “incompatible” *V. dahliae* strains under starvation

Starvation induces significantly CAT-mediated fusion of *V. dahliae* conidia/germlings (Vangalis *et al*., 2021), and this motivated us to investigate the possibility of conidial fusion between strains of different VCGs under such conditions. Seven representative strains (Table S1; Fig. 1a) were labeled with sGFP- or mCherry-tagged histone H1 (nuclear localization) for their microscopic identification in pairings and, therefore, the discrimination of fusion between identical or unlike cells. In 14 pairings of the labeled strains, inter-strain fusion (Fig. 1b) was observed at significant levels (5.2-33.8% of fusions) not only in “compatible” pairings (i.e. same VCG), but also in all “incompatible” combinations (i.e. different VCGs) (no significant difference between “compatible” and “incompatible” pairings; Student’s *t*-test, *p*-value > 0.05; Fig. 1c). These results demonstrate that spontaneous cell fusion occurs frequently under starvation conditions between conidia or germlings of different strains, regardless of their previous VCG classification.

**Fig. 1.**
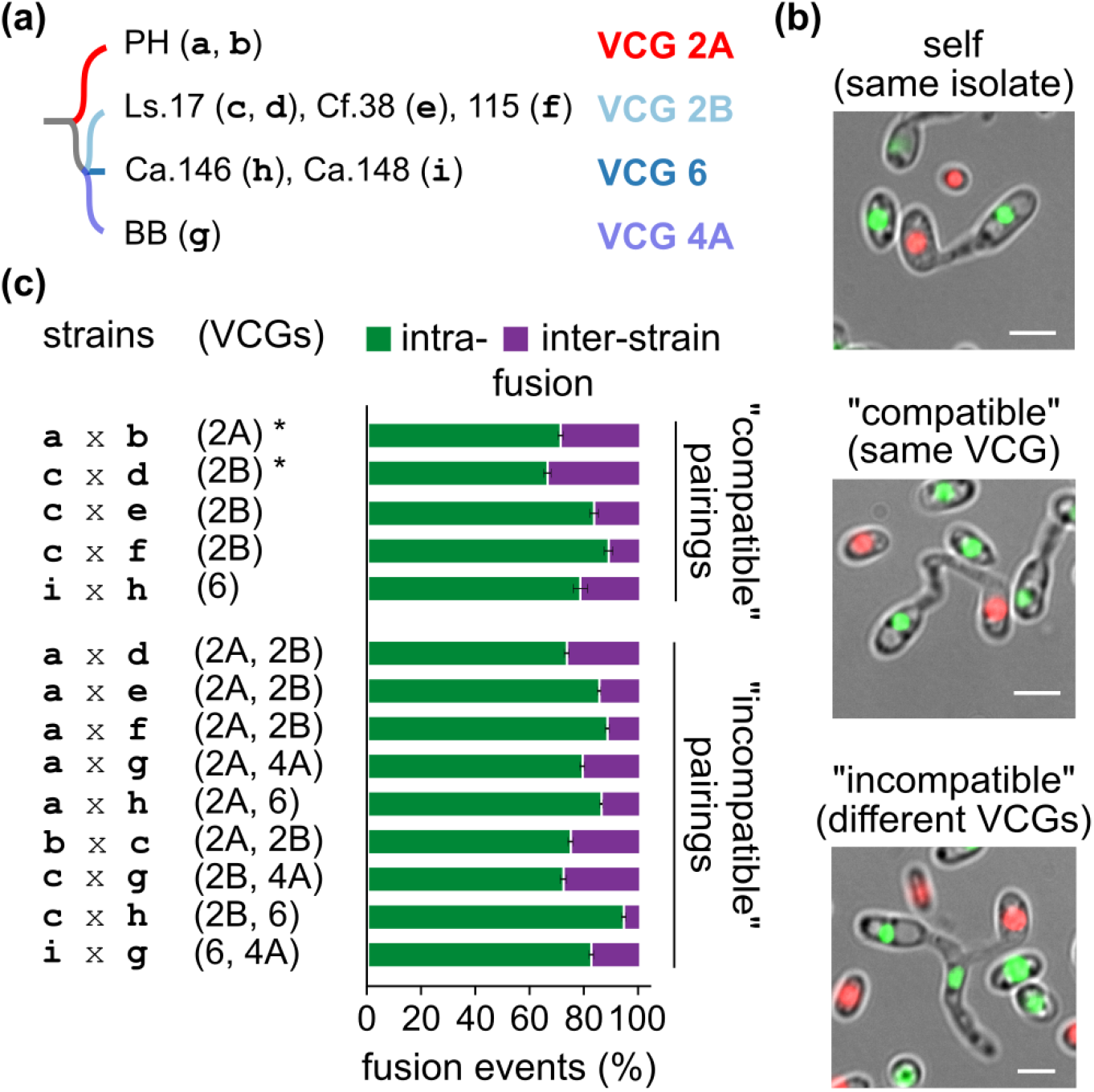
Spontaneous non-self fusion via CATs occurs frequently between “incompatible” strains of *V. dahliae* under starvation. (a) Wild-type strains used in this experiment and their VCG classification. Single-letter strain codes (in brackets) denote derivative strains expressing sGFP- or mCherry-tagged histone H1 (details in Table S1). The structure of the dendrogram reflects the phylogenetic relationships of VCGs as these were previously determined (Papaioannou *et al*., 2013a). (b) Examples of CAT-mediated fusion between conidia/germlings of strains c-d (originating from the same wild-type strain), c-e (members of VCG 2B) and c-g (“incompatible” members of VCGs 2B and 4A, respectively). Bars = 5 μm. (c) Frequencies of intra- and inter-strain fusion in pairings of *V. dahliae* strains a-i. Control combinations indicated with an asterisk involved strains originating from the same wild-type strain. Each pairing was tested in triplicate, and 250-300 fusion events were analyzed per replicate. Bars = SD.

### Heterokaryotic cells arisen from CAT-mediated fusion of “incompatible” strains often escape incompatibility-triggered cell death

We next asked whether fusion of “incompatible” conidia/germlings could lead to the establishment of viable heterokaryotic cells. To investigate this, we paired labeled strains (expressing fluorescently tagged histone H1 and/or cytoplasmic sGFP; Table S1) and used live-cell imaging to monitor the cell fate and nuclear behavior in self, “compatible” and “incompatible” fusions over time, up to 24 h after fusion. In total, we collected time-course data from 908 individual fusions, from three self (304 events), three “compatible” (80 events) and four “incompatible” (524 events) pairings (Table S5). Overall, in this time window we identified four distinct types of cellular behavior following CAT-mediated fusion (Fig. 2a,b):

**Fig. 2.**
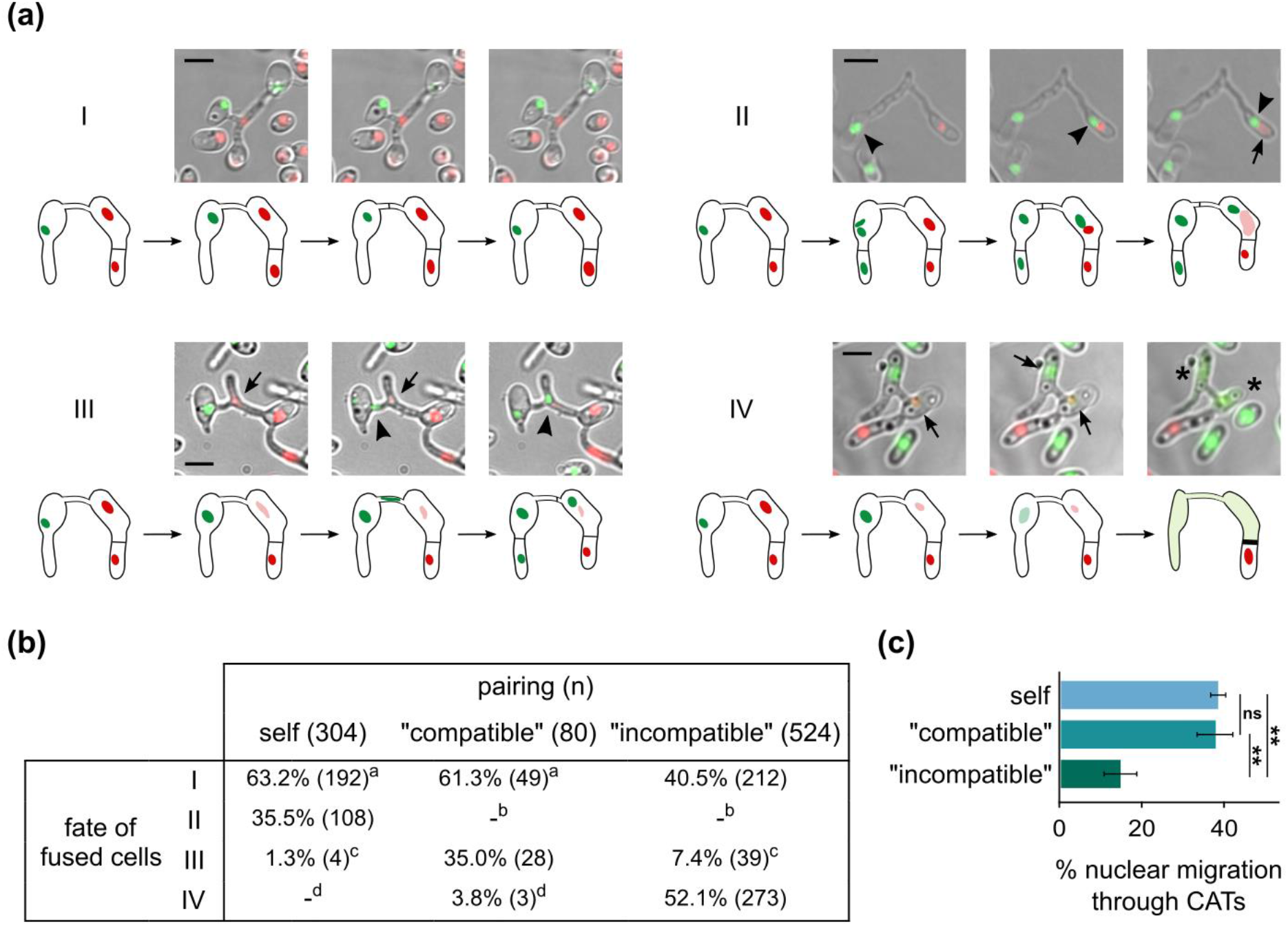
“Incompatible” fused cells (via CATs) often escape incompatibility-triggered cell death. (a) Examples of the four observed types of cellular behavior following CAT-mediated fusion of “compatible” and “incompatible” cells. Strains shown: Ls.17 H1-mCherry paired with Cf.38 H1-sGFP (I), Ls.17 H1-sGFP (II), PH H1-sGFP (III) and BB H1-sGFP (IV). Arrows: nuclei undergoing degradation; arrowheads: migrating nuclei; asterisks: cell shrinkage. Bars = 5 μm. (b) Frequencies of the four types of cellular behavior in self, “compatible” and “incompatible” pairings. The numbers of detected events in each case are provided in brackets (detailed results in Table S5). (c) Frequency of nuclear migration through CATs in viable heterokaryons resulting from self, “compatible” and “incompatible” pairings. Bars = SD. In (b,c): statistical significance of differences between the compared groups was tested with one-way ANOVA followed by Tukey’s post-hoc test; in (b): groups marked by the same superscript character did not differ significantly; in (c): ** *p*-value ≤ 0.01; ns: non-significant (*p*-value > 0.05).

I. Fusion led to a viable anastomosed cell. No nuclear migration occurred through the CAT, and septae were frequently formed within the CAT (70%). This behavior was frequently observed in self and “compatible” pairings (62.3%), but also in “incompatible” combinations (40.5%). In “compatible” pairings, cytoplasmic intermixing between the fused cells was always observed, whereas that was the case in 63.6% of “incompatible” cases (Movie S1).

II. Fusion was followed by a nuclear division in one of the interacting cells (20-150 min after fusion). One of the daughter nuclei then migrated through the CAT from the donor cell to its fusion partner (migration was completed in 10-20 min). Following a dikaryotic phase in the recipient cell (10 min -3 h), its resident (i.e. original) nucleus was degraded. This type was only detected in self-pairings (35.5%).

III. Similar to (II), but degradation of the resident nucleus happened before migration of the other nucleus. This was often observed in “compatible” pairings (35.0%), less frequently in “incompatible” (7.4%), and only rarely in self-interactions. Nuclear migration through a CAT (types II and III) was often (59.3%) followed by formation of a septum in the CAT, regardless of the “compatibility” classification of the paired strains.

IV. Fusion was followed by a cell death reaction observed in half of the “incompatible” pairings tested, and rarely in “compatible” interactions. The catastrophic reaction started 10-120 min after fusion, with or without obvious cytoplasmic mixing (Fig. S1a,b). It was generally restricted to the fused cellular compartments without affecting the adjacent cells, and it typically involved thickening of the surrounding septae and cell walls, degradation of both nuclei and gradual cell shrinkage (Movie S2; Fig. S1c).

Fatal incompatibility reactions following CAT-mediated fusion were essentially limited to “incompatible” fusions but affected only half of them. The considerable remaining fraction (47.9%) apparently escaped cell death and formed stable heterokaryotic cells without any signs of cellular degeneration in the analyzed time window, in all tested combinations of strains (Table S5). Cell viability was confirmed by the addition at the end of each experiment of methylene blue, which generally fails to stain viable heterokaryons, in contrast to dead cells (Fig. S1d). Nuclear translocation through the CAT was detected at significant levels in all types of pairings, including “incompatible” interactions (Fig. 2c). These results demonstrate that fusion of “incompatible” conidia/germlings can lead to the establishment of viable heterokaryons with the possibility of nuclear migration through their anastomosis bridge, at significant frequencies. Nuclear translocation is invariably associated with selective degradation of the nucleus of the recipient cell, both in “compatible” and “incompatible” fusions (Fig. 2b).

### “Incompatible” fusions that escape cell death support heterokaryotic growth in colonies

Since CAT-mediated fusion of “incompatible” strains can often lead to viable heterokaryotic cells, we hypothesized that these could further support hyphal growth to the formation of heterokaryotic colonies. We tested this using a collection of 19 nitrate non-utilizing (*nit*) strains of all *V. dahliae* VCGs, as well as two strains of the related species *V. nonalfalfae* (Table S1). When hyphae of “incompatible” complementary *nit* strains are confronted on minimal medium (MM) that selects for prototrophy, typically no heterokaryotic growth occurs due to incompatibility barriers (Fig. 3a). We performed 110 pairings using this hyphal-based traditional assay (Fig. 3a), in comparison to an assay that permitted CAT-mediated cell fusion of the same pairs of strains (Fig. 3c). The results were drastically different (Fig. 3b,d; Table S6), with the CAT-based method yielding 5.5 times more positive results than the hyphal-based assay, in combinations of all VCGs. In total, 49.0% of the “incompatible” pairings tested positive only in the CAT-based assay. The latter yielded several interspecific responses between *V. dahliae* and *V. nonalfalfae*, as well as interactions involving a heterokaryon self-incompatible (HSI) strain (Fig. 3d).

**Fig. 3.**
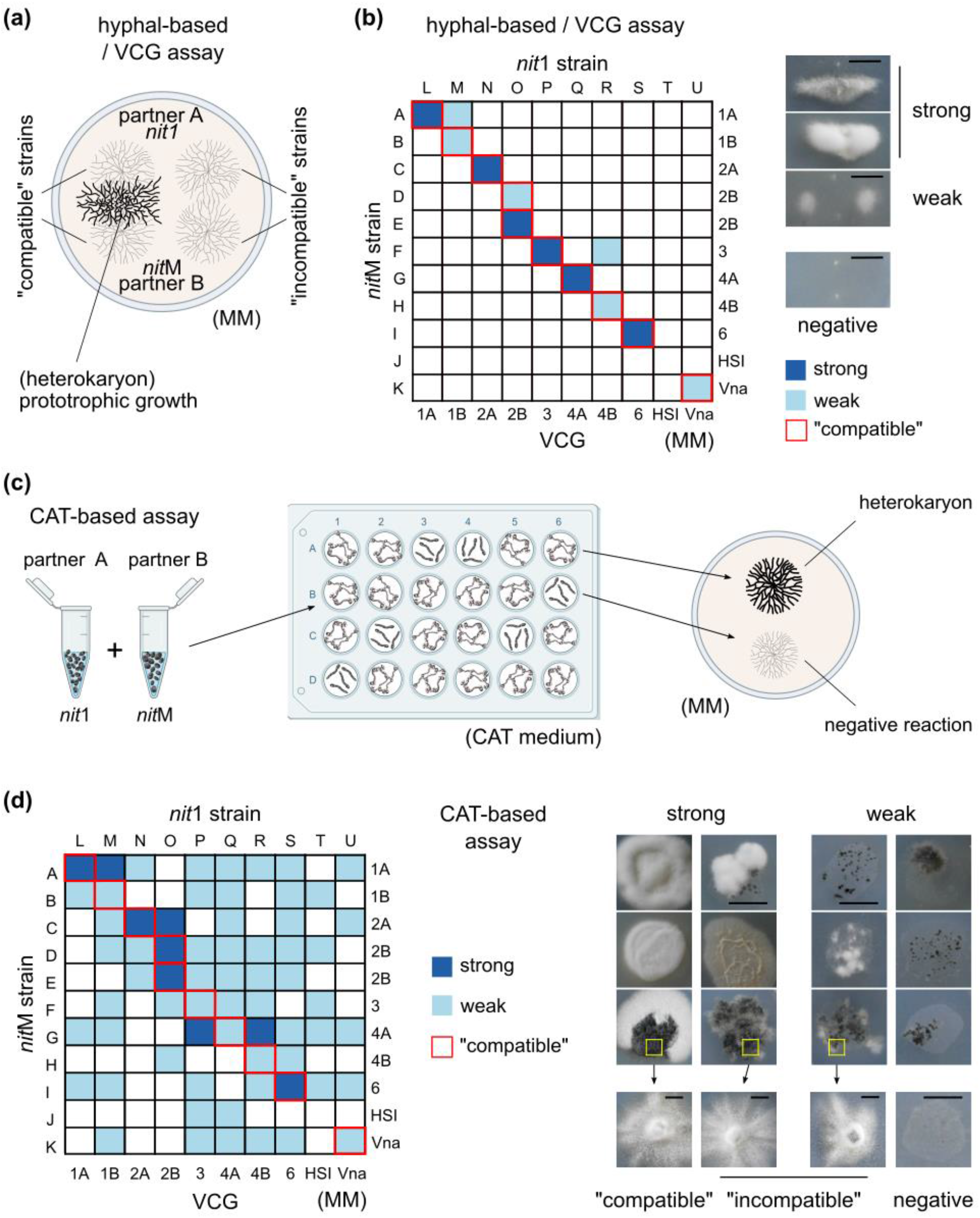
Conidial/germling fusion (via CATs) supports stable growth of “incompatible” heterokaryotic colonies. The traditional (hyphal-based) VCG (a) and the new CAT-based (c) complementation assays for the detection of heterokaryotic growth are described schematically. The results from the two assays are summarized in (b) and (d), respectively (see Table S6 for details). A single-letter code has been assigned to each strain (details of strains are provided in Table S1). HSI: heterokaryon self-incompatible. Vna: *V. nonalfalfae*. Each pairing (in both assays) was performed in at least three replicates. Pairings that produced very limited prototrophic growth in only one of the replicates were regarded as negative. Examples of strong and weak heterokaryons (21 days old) are shown for each method (right), as well as their morphological characteristics after subculturing them on fresh MM (arrows). Bars = 2 cm.

Most “incompatible” interactions were weak, exhibiting sparse aerial mycelium, tufts of hyphae, or localized groups of microsclerotia in at least two out of the three replicates, as opposed to the strong interactions with denser prototrophic growth in all three replicates (Fig. 3b,d). We sought, therefore, to validate the heterokaryotic nature of such colonies by excluding the possibilities of reversion to prototrophy or cross-feeding of non-fused cells. When 30 randomly selected weak “incompatible” heterokaryons were analyzed, all retained the ability to grow slowly over successive reculturings on fresh MM, with denser growth in their central regions, irregular colony shape with frequent morphological sectoring, and uneven pigment distribution, which are all typical features of heterokaryons (Fig. 3d). We then plated approx. 10^4^ uninucleate conidia of each heterokaryon on MM and we never detected prototrophic growth, but only *nit*-characteristic sparse growth. These results confirmed the heterokaryotic nature of the hyphal masses. Our experiments suggest that large numbers of fusions occur even between “incompatible” strains and that sufficiently many of them escape incompatibility-triggered cell death to support prototrophic growth of the resulting colonies.

### Heterokaryosis arises from mixed infections *in planta*

With the exception of its dormant microsclerotia in the soil, *V. dahliae* completes its life cycle in its plant host (Pegg & Brady, 2002). To examine whether the parasexual cycle in nature is initiated by heterokaryon formation within plants, we subjected eggplant seedlings to mixed artificial infections (Fig. 4a) with “compatible” (Ls.17 × Cf.38) and “incompatible” (Ls.17 × PH; Ls.17 × BB; Ls.17 × Ca.146) combinations of strains, which were marked with complementary antibiotic-resistance markers (Table S1). Fungal material was re-isolated from stem sections of the infected plants 40 days after inoculation. Presumably heterokaryotic growth was detected on double-selective PDA medium (amended with both antibiotics) in 16 out of 425 tested xylem chips (3.8%), from all “compatible” and “incompatible” combinations, whereas no growth was observed in any of our single-infection or mock-infection controls (Fig. 4b,c). All 16 strains retained their ability to grow slowly over successive reculturings on double-selective medium, but no growth was detected when approx. 10^4^ uninucleate conidia of each sample were plated on that medium, thus confirming the heterokaryotic nature of the 16 samples. Given the complete suppression of conidial fusion on PDA (Vangalis *et al*., 2021), our results suggest that the isolated “compatible” and “incompatible” heterokaryons arose from fusion between strains *in planta*.

**Fig. 4.**
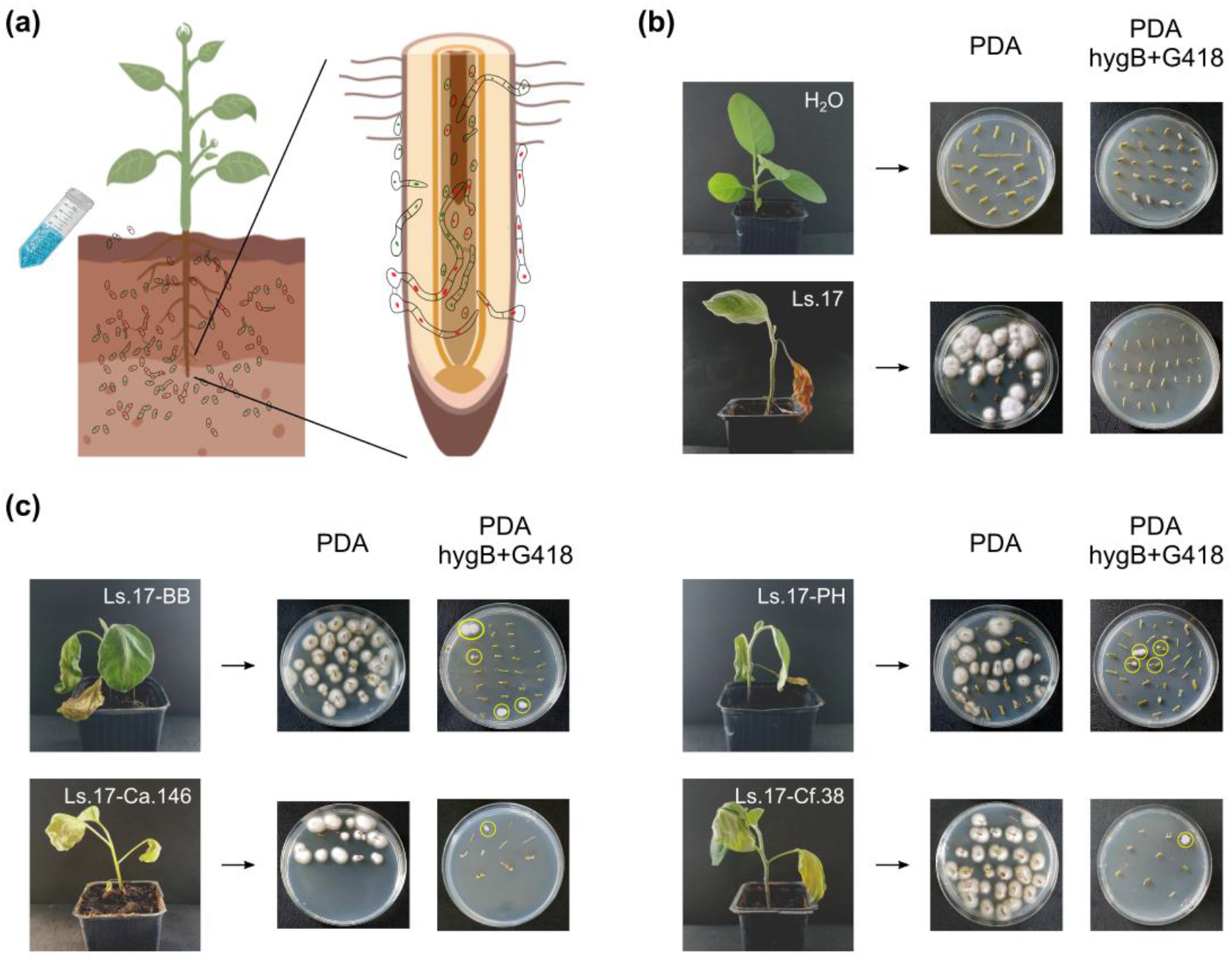
Heterokaryon formation *in planta* following mixed infection. (a,b) Eggplant seedlings were artificially infected with mixed inocula of “compatible” and “incompatible” combinations of *V. dahliae* strains. The pathogen was re-isolated from stem sections (from eight xylem chips along the stem of each plant) of 17 infected plants per strain combination, 40 days after inoculation. From each treated plant, three xylem chips were transferred to PDA (controls of re-isolation), while the remaining five were transferred to double-selective PDA supplemented with hygB and G418, to select for heterokaryotic growth. Mock-infected plants (with water) and single-strain infections were used as controls. (c) Similarly to (b), for seedlings infected with mixed inocula of strains. “Compatible” pairing: Ls.17 × Cf.38; “incompatible” pairings: Ls.17 × PH, Ls.17 × BB, Ls.17 × Ca.146.

### Optimization of an efficient CRISPR/Cas9-mediated gene targeting system for the analysis of autophagy in *V. dahliae* heterokaryons

The characteristics of incompatibility-triggered cell death and selective nuclear degradation following cell fusion (Figs 2, S1) indicated that the autophagic machinery of the cells may be an important component of the mechanisms that regulate the heterokaryotic cell fate decision and nuclear interaction. To investigate this, we first searched the *V. dahliae* genome for homologs of the core autophagy genes and discovered that *V. dahliae* possesses a full autophagy gene complement (Table S3). We selected the hallmark autophagy genes *atg1* (VDAG_05745) and *atg8* (VDAG_01225), which are essential for the induction of autophagy and formation of autophagosomes, respectively (Feng *et al*., 2014), to investigate the possible roles of autophagy in the post-fusion processes.

To overcome practical limitations that complicate reverse genetic investigations in *V. dahliae*, we used the AMA1 replicator sequence from *Aspergillus nidulans* (Gems *et al*., 1991) to ensure autonomous plasmid maintenance in a CRISPR/Cas9-based system (Nødvig *et al*., 2015) for efficient gene targeting in *V. dahliae*. For plasmid delivery to the fungal cells we developed a lithium acetate/heat-shock-mediated method for conidial transformation. This technique yielded lower transformation efficiencies than our standard protoplast-based method (up to 17 and 106 transformants μg^-1^ of plasmid, respectively), but it proved much faster and more convenient. Using this method, we transformed *V. dahliae* Ls.17 with a plasmid containing the hygromycin B-resistance (*hph*) selection marker, and we studied plasmid stability over 14 successive reculturings of 25 randomly selected transformants. While full stability was observed on selective medium, a gradual decrease in plasmid stability occurred in the absence of selection (Fig. S2a). This pattern proved very convenient, as it offered clear-cut selection of transformants and maintenance of strains under selective conditions, but also straightforward elimination of the plasmid containing the *Spcas9* endonuclease gene and the selection marker, after the targeted genetic modification had been achieved.

We then constructed an AMA1-containing plasmid for the CRISPR/Cas9-mediated targeting of the N-terminal region of *V. dahliae atg8* (Fig. 5a). In this plasmid, the constitutive P*gpdA* promoter was used for the pol-II-mediated transcription of the sgRNA cassette. The sgRNA was flanked by the ribozymes HH and HDV, which ensure the post-transcriptional release of the sgRNA without modifications (Fig. 5a). The plasmid was used to co-transform V. *dahliae* Ls.17 together with a linear repair substrate consisting of either the *sgfp* or the *mCherry* coding sequence, flanked by homologous arms corresponding to the genomic regions that surround the start codon of *atg8* (Fig. 5b). Using this method, we achieved *in situ* seamless (i.e. without introducing any additional sequence) marker-free tagging of *atg8* (Fig. 5c), with efficiencies that ranged between 12-66% of transformants for 100-1,000 bp-long homologous flanks, respectively (Fig. S2b). Properly tagged strains were validated using PCR and microscopic analyses that confirmed the induction of autophagy by starvation and treatment with rapamycin, which is a known inducer of autophagy (Mizushima, 2007) (Fig. S2c), as well as the previously described involvement of autophagy in conidial germination (Kikuma *et al*., 2006) (Fig. S2d).

Next, we used the same system for inactivation of *V. dahliae atg8* by gene disruption. For this, the G418-resistance (*neo*^R^) cassette, flanked by 100 bp-long homologous arms (Fig. 5b), was used for co-transformation of *V. dahliae* Ls.17 with the plasmid containing the sgRNA cassette (Fig. 5a). To ensure inactivation of the gene following homologous recombination, stop codons were introduced in the arm downstream of the *neo*^R^ cassette (Fig. 5c). This approach for gene knockout proved faster than the commonly used *Agrobacterium tumefaciens*-based system (ATMT) (Fig. S3a), which we used to delete *atg1*. In both cases, knockout mutants were verified by PCR and Southern hybridization analyses (Fig. S3b,c). The CRISPR/Cas9-based system yielded correctly disrupted mutants at a relatively low frequency (1.1% of transformants) using 100 bp-long arms, which conveniently overcomes the need for multistep vector construction.

**Fig. 5.**
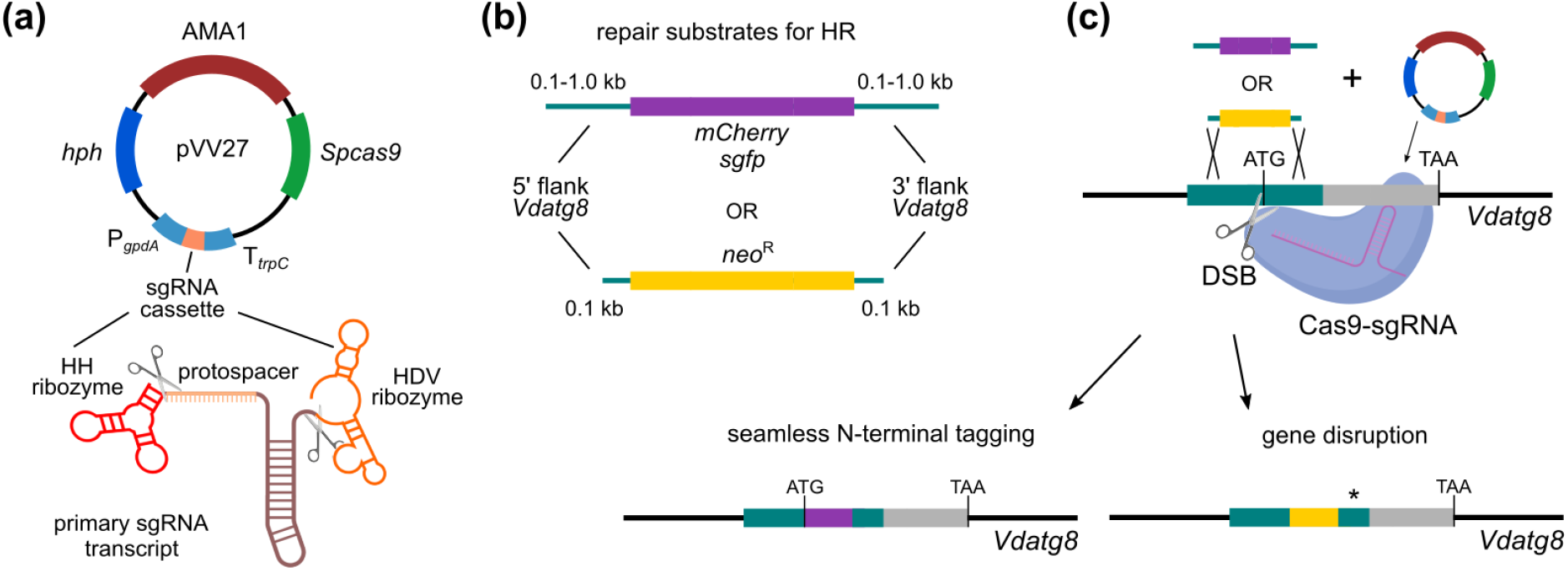
Schematic representation of CRISPR/Cas9-mediated gene targeting in *V. dahliae*. (a) Both SpCas9 and sgRNA cassettes were introduced into the same, AMA1-containing plasmid. The sgRNA was flanked by the hammerhead (HH) and hepatitis delta virus (HDV) ribozymes, which enable the release of the mature sgRNA after their self-catalysis (scissors indicate cleavage points). (b) Homologous repair substrates consist of either a fluorescent protein gene (for gene tagging) or an antibiotic-resistance cassette (for gene disruption), flanked by homologous arms of variable length. (c) Co-transformation of the AMA1-containing plasmid with the homologous repair substrate for gene tagging or disruption. The Cas9/sgRNA complex induces in the genomic locus targeted DSBs (scissors), which can be repaired using the homologous repair substrate to result in seamless gene tagging or gene disruption. In the latter case, stop codons (asterisk) were introduced in the homologous arm downstream of the antibiotic-resistance cassette, to ensure the knockout of the gene (example shown: *V. dahliae atg8*).

### Autophagy is involved in post-fusion selective nuclear degradation but is not required for cell fusion and acts against incompatibility-triggered cell death

In order to gain further insight into the roles of autophagy in heterokaryosis, we first constructed a *V. dahliae* H1-mCherry sGFP-Atg8 strain (Ls.17 background), which we used for time-lapse imaging of CAT-mediated self-fusion. Following anastomosis, we invariably observed co-localization of nuclei undergoing selective degradation (Figs 2a, S1c, 6a,b, S2e; Movie S3) with Atg8-containing organelles. These were ranging from small autophagosomes associated with parts of the nuclear chromatin (Fig. S4) to larger and sometimes ring-like structures that appeared to engulf the whole nucleus before its degradation (Fig. 6a; Movie S3). During the Atg8-nuclear interaction, the Atg8-containing vesicles were often dynamically undergoing fusion and fission (Fig. S4).

**Fig. 6.**
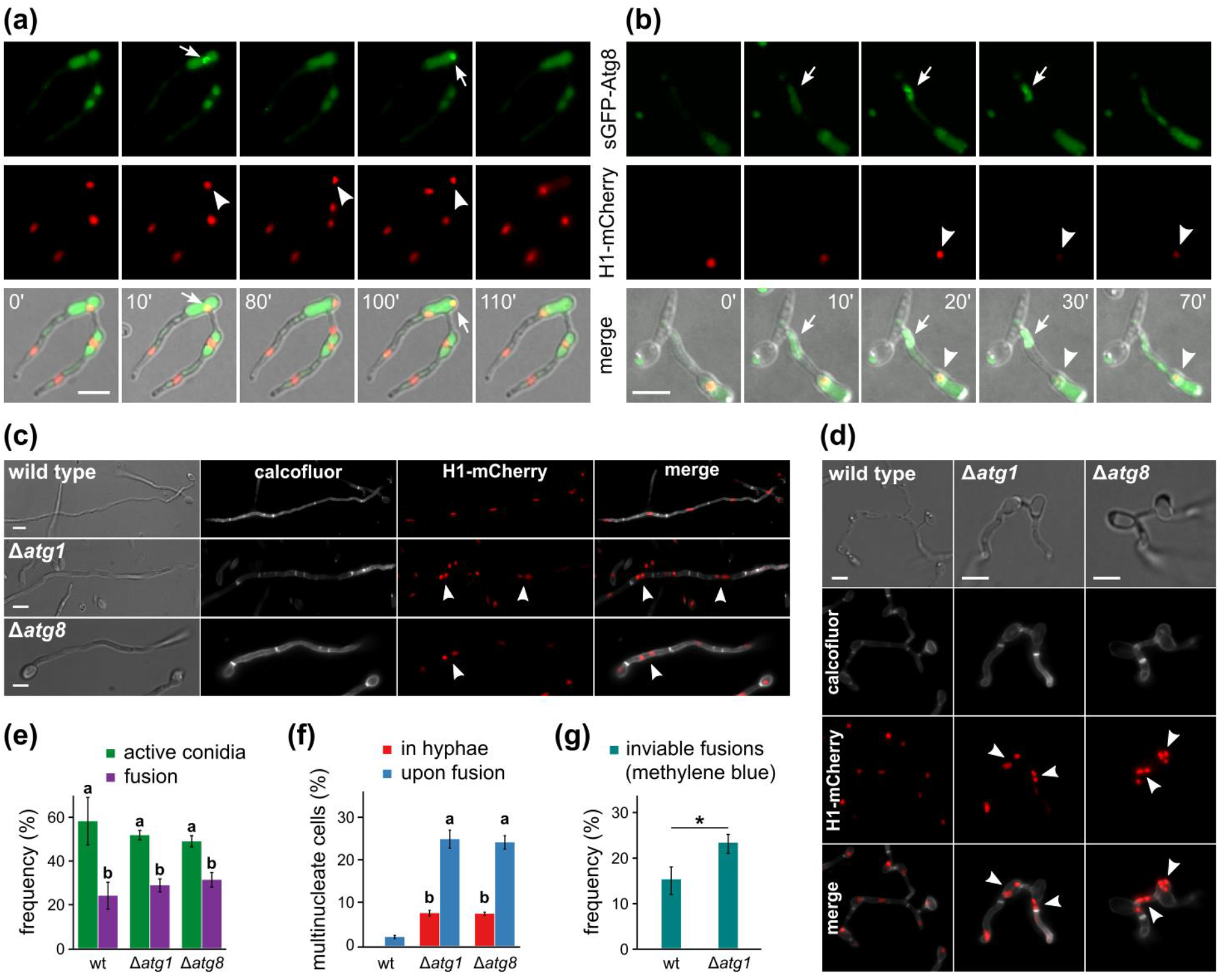
Autophagy is involved in selective nuclear degradation in heterokaryons, but it is not required for fusion or cell death due to incompatibility. (a) Live-cell imaging of a self-fusion (Ls.17 H1-mCherry sGFP-Atg8). A ring-like structure with accumulated sGFP-Atg8 (arrows) surrounds the sequestered nucleus (arrowheads). (b) Live-cell imaging of an “incompatible” fusion (Ls.17 H1-mCherry sGFP-Atg8 × BB H1-sGFP). Localized accumulation of sGFP-Atg8 near the fusion point (arrows) shortly before the incompatibility reaction. Arrowheads: nucleus undergoing degradation. (c,d) Multinucleate cells (arrowheads) arise frequently in Δ*atg1* and Δ*atg8* autophagy-deficient mutants from sub-apical nuclear division (c) or cell fusion (d), in contrast to the wild type that exhibits strictly uninucleate organization. Calcofluor white was used for cell wall staining. Bars in (a-d) = 5 μm. (e) Frequency of active conidia and their fraction involved in CAT-mediated fusion. Each strain was tested in triplicate (n = 300 conidia per replicate). (f) Frequency of non-apical hyphal compartments and fused cells with more than one nuclei. Each strain was tested in triplicate (n = 300 hyphal cells or 150 anastomoses per replicate). In (e,f): statistical significance of differences was tested with one-way ANOVA followed by Tukey’s post-hoc test; bars with the same letter do not differ significantly (*p*-value > 0.05). (g) Frequency of inviable fusions, as determined by staining with methylene blue. Wild-type (wt) pairing: Ls.17 H1-mCherry × PH H1-sGFP; pairing of autophagy-deficient mutants (Δ*atg1*): Ls.17 Δ*atg1* × PH Δ*atg1*. Each pairing was tested in triplicate (n = 150 anastomoses per replicate). Statistical significance of difference between the two groups was assessed using the Student’s *t*-test (* *p* ≤ 0.05). Error bars in (e-g): SD.

In our autophagy-deficient mutants Δ*atg1* and Δ*atg8*, conidial germination and CAT-mediated fusion occurred at wild-type levels (Fig. 6e), in contrast to *F. oxysporum* where deletion of *atg8* was previously reported to impair its fusion frequency (Corral-Ramos *et al*., 2015). Both *V. dahliae* mutants exhibited significantly more bi- and multi-nucleate fusions than the wild type (Fig. 6d,f; Movie S4), whereas gene complementation restored the wild-type phenotype in both cases (Fig. S5). Similarly, bi-/multi-nucleate cells arose frequently from nuclear division in Δ*atg1* and Δ*atg8* hyphal compartments (Fig. 6c,f; Movie S5), in contrast to the strictly uninucleate organization of the wild-type mycelium (Fig. 6c,f), which is ensured by degradation of one of the daughter nuclei of each non-apical nuclear division (Movie S6). Overall, our results suggest that the nuclear number is strictly regulated in *V. dahliae* to one per cell, and this process is mediated by the autophagic machinery.

We then tested whether autophagy is involved in the post-fusion incompatibility-triggered reaction, which displays morphological features reminiscent of autophagic programmed cell death (Figs 2a, S1). Time-lapse imaging of “incompatible” fusions between strains Ls.17 H1-mCherry sGFP-Atg8 and BB H1-sGFP revealed in all cases localized accumulation of Atg8 in the vicinity of the fusion point, prior to the onset of the catastrophic reaction (Fig. 6b). Co-localization of nuclei undergoing degradation with Atg8-containing vacuoles was also observed in all “incompatible” fusions (Fig. 6b). To investigate whether autophagy mediates the incompatibility reaction, we also deleted *atg1* from strain PH and analyzed a PH Δ*atg1* × Ls.17 Δ*atg1* pairing. Staining of inviable fusions with methylene blue (Fig. S1d) revealed a significantly higher frequency of cell death in the autophagy-deficient pair than in the wild-type pair (Fig. 6g), while typical morphological characteristics of incompatibility were observed in affected heterokaryons in both cases. Therefore, although autophagy is induced before the manifestation of the incompatibility reaction, it is not required for cell death but actually protects some “incompatible” fusions from destruction. This is in line with previous studies of *P. anserina* that proposed a possibly protective role of autophagy against incompatibility-triggered cell death (Pinan-Lucarré *et al*., 2003, 2005).

### Nuclear interactions in “compatible” and “incompatible” heterokaryons

Co-existence of genetically distinct nuclei in heterokaryons induces the parasexual cycle and may enable horizontal gene transfer, but the mechanisms involved in these nuclear interactions are not well understood. Despite our many attempts, we were unable to capture direct fusion of nuclei in our heterokaryons, which is explained by the very low frequencies of diploidization (10^-6^-10^-8^) in *V. dahliae* heterokaryons (Typas & Heale, 1978). However, in a few cases we observed gradual accumulation of sGFP signal in mCherry-labeled nuclei or *vice versa*, both in “compatible” and in “incompatible” pairings (Fig. 7a). In these cases, transfer was strictly unidirectional and not always correlated temporally with nuclear division. Although the underlying mechanism is unknown, such interactions provide evidence for genetic transfer between nuclei. In addition, we analyzed CAT-mediated self-fusions of Ls.17 Δ*atg1* cells to examine whether their higher frequencies of binucleate cells facilitate the microscopic study of nuclear fusion (Fig. 6f). Indeed, fusion of nuclei was detected in rare cases in this strain (Fig. 7b).

**Fig. 7.**
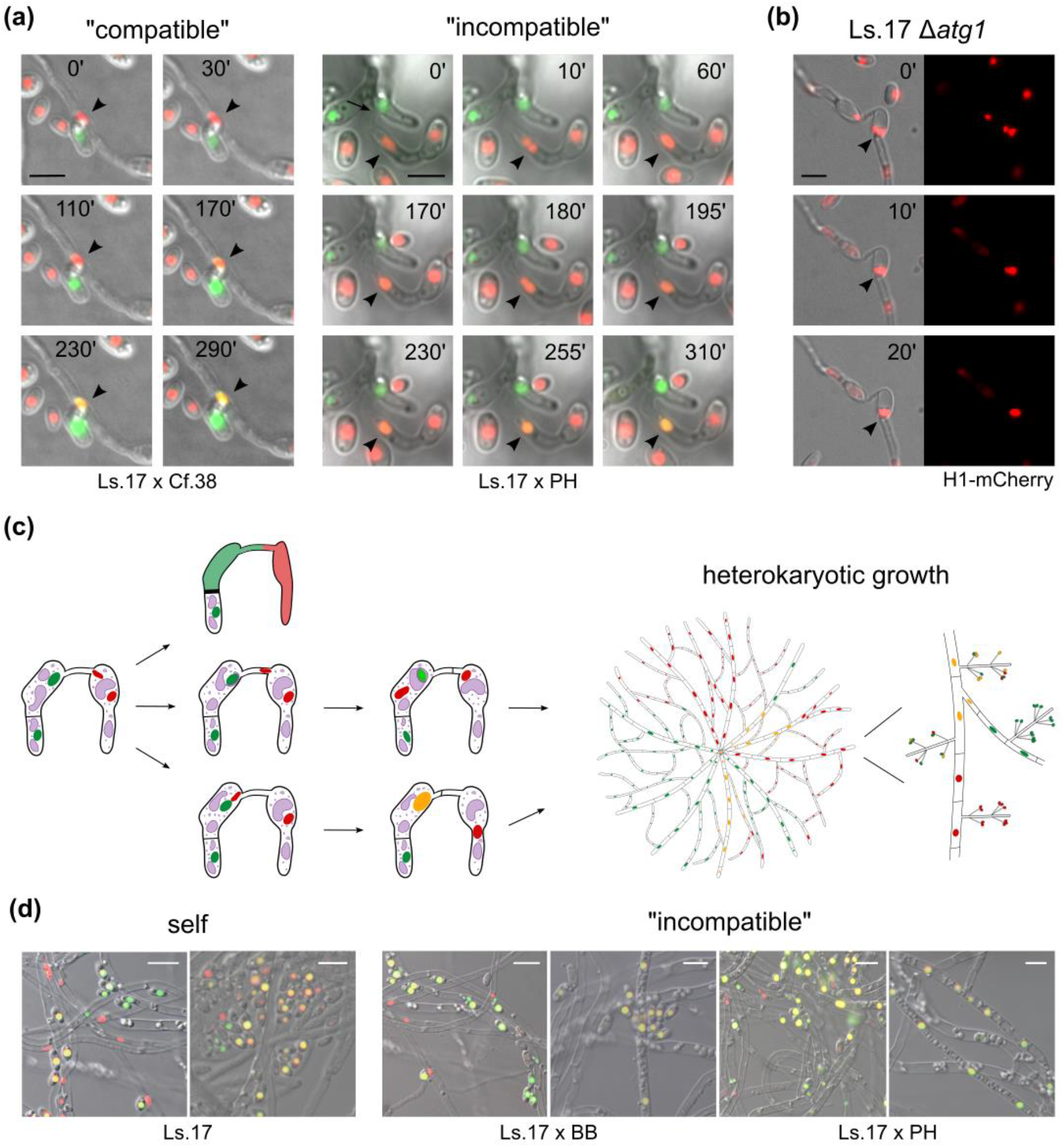
Nuclear interactions in “compatible” and “incompatible” *V. dahliae* heterokaryons. (a) Gradual accumulation of sGFP signal in mCherry-labeled nuclei in heterokaryotic cells between strains Ls.17 H1-mCherry × Cf.38 H1-sGFP (“compatible”), and Ls.17 H1-mCherry × PH H1-sGFP (“incompatible”). Arrowheads: recipient nuclei. (b) Direct nuclear fusion (arrowhead) in strain Ls.17 Δ*atg1*. Bars in (a,b) = 5 μm. (c) Schematic representation of the possible fates of a non-self cell fusion in *V. dahliae*. Fused cells can remain viable or be subjected to an incompatibility-triggered reaction. Division of rare fused nuclei in viable heterokaryotic cells can generate distinct nuclear lineages in mosaic colonies. (d) Nuclei of “incompatible” (Ls.17 H1-mCherry × BB sGFP, Ls.17 H1-mCherry × PH sGFP) and self-pairing control (Ls.17 H1-mCherry × Ls.17 sGFP) heterokaryons following growth under non-selective conditions. Bars = 10 μm.

As long as they are able to divide, fused nuclei can form distinct nuclear lineages in chimeric heterokaryotic colonies (Fig. 7c). To investigate whether this is possible in “incompatible” interactions, we scanned microscopically heterokaryotic colonies formed via CATs, under non-selective conditions, between strains with sGFP- or mCherry-labeled nuclei. Both our self-pairing control (Ls.17) and the “incompatible” fusions (Ls.17 × BB, Ls.17 × PH) gave rise to mosaic colonies with mixed populations of nuclei with sGFP, mCherry or both signals (Fig. 5d). These colonies mainly consisted of regions with parental properties, interspersed with sectors or individual hyphae (and uninucleate conidia) with mixed fluorescent signal, which is consistent with the mosaic nature of *V. dahliae* heterokaryons described earlier (Puhalla & Mayfield, 1974). Our results indicate that parasexual interactions are possible in “incompatible” heterokaryons of *V. dahliae*.

## Discussion

Despite the evolutionary benefits and persistence of sexual reproduction across the eukaryotic tree of life (Muller, 1932; Felsenstein, 1974; Otto & Lenormand, 2002), loss of sexual competence is observed in members of several eukaryotic lineages, including approx. one fifth of described fungi (i.e. the mitosporic fungi) that either appear to be strictly asexual or are suspected to engage only rarely in some cryptic form of sex (Taylor *et al*., 1999; Dyer & Kück, 2017). These organisms are very important to plant pathology and their effective management depends on our ability to predict successfully the modes and pace of their evolution. Central questions in this regard are what are the relative contributions, mechanisms and spatiotemporal patterns of clonal reproduction and recombination in these organisms (Taylor *et al*., 1999). Here, we used *V. dahliae*, an asexual pathogen, to gain insights into the extent and mechanisms of heterokaryosis, an essential component of parasexuality, which enables genomic recombination in the absence of sex and meiosis (Pontecorvo, 1956; Clutterbuck, 1996; Bennett, 2015; Strom & Bushley, 2016). We found that starvation induces fusion between conidia or germlings of different strains, regardless of their traditionally assessed vegetative compatibility (Leslie, 1993). Cell fusion can often lead to the establishment of viable heterokaryons, where nuclei can interact to initiate the parasexual cycle.

The VCGs of asexual fungi, assessed by forcing hyphae of complementary auxotrophic mutants to anastomose on minimal medium, have been widely regarded as incompatible groups (Puhalla, 1985; Leslie, 1993). However, cross (or “bridging”) inter-VCG reactions have been reported in some species (Joaquim & Rowe, 1990; Molnár *et al*., 1990; Kerényi *et al*., 1997; Viji *et al*., 2004; Papaioannou *et al*., 2013b), and their genetic analysis in *V. dahliae* previously revealed that such interactions can be due to heterokaryosis (Papaioannou & Typas, 2015). In addition, studies of CAT-mediated fusion of *Colletotrichum lindemuthianum* (Ishikawa *et al*., 2012) and *Fusarium oxysporum* (Shahi *et al*., 2016) suggested that incompatibility may be suppressed at the conidial/germling stage of development. However, our previous investigation of conidial pairings of *V. dahliae* strains in minimal medium showed excellent congruence with the traditional hyphal pairings on agar medium (Papaioannou & Typas, 2015), providing no evidence of incompatibility suppression during conidial germination in *V. dahliae*.

We recently discovered that starvation of cells induces significantly CAT-mediated fusion of *V. dahliae* conidia/germlings (Vangalis *et al*., 2021). Here, we show that this happens regardless of their VCG classification, even between species, and that inter-VCG fusions often form viable heterokaryotic cells and colonies. No selection pressure for heterokaryons was exercised in our experiments, in sharp contrast to the traditional VCG assays that force heterokaryosis between auxotrophic strains under non-physiological selection conditions. In addition, we demonstrated that heterokaryosis can result from mixed infections *in planta*, which constitute the most likely opportunity for encounter of non-dormant *V. dahliae* strains in nature (Wheeler & Johnson, 2019). Overall, our study demonstrates that the traditionally assessed incompatibility systems do not constitute absolute barriers to heterokaryon formation in *V. dahliae*, where a fraction of starved conidia can escape allorecognition systems, anastomose and lead to the formation of viable heterokaryons, regardless of their VCG classification. Our preliminary experiments revealed that starvation similarly induces anastomosis between mature hyphae of strains on water agar. A presumed extension of the imperfect nature of incompatibility systems, observed here in fusion of conidia, to hyphal fusion could explain the frequently observed cross-VCG hyphal interactions in this species (Joaquim & Rowe, 1990; Elena & Paplomatas, 1998; Hiemstra & Rataj-Guranowska, 2003; El-Bebany *et al*., 2013; Papaioannou *et al*., 2013b). It would also suggest that it is the environmental conditions (e.g. starvation) rather than the developmental stage that induces anastomosis and parasexual phenomena even between previously considered “incompatible” strains.

Multiple checkpoint systems have evolved in the sexual ascomycete *N. crassa* to essentially prevent heterokaryosis between genetically distinct individuals (Daskalov *et al*., 2017; Gonçalves & Glass, 2020). Our study indicates that partner recognition and chemotropic attraction preceding cell fusion in *V. dahliae* are not subject to the same genetic control as in *N. crassa*, where an allorecognition system prevents fusion between incompatible individuals (Heller *et al*., 2016). Nevertheless, we found evidence for the existence of two downstream allorecognition checkpoint systems following cell fusion. First, a fraction of “incompatible” interactions failed to achieve cytoplasmic mixing, which may have been prevented by a system similar to that of *N. crassa*, where allelic differences between individuals at the *cwr* locus can cause cell fusion arrest by blocking dissolution of the cell wall (Gonçalves *et al*., 2019). Secondly, in a fraction of “incompatible” interactions fusion triggered a cell death reaction characterized by hyphal compartmentalization, thickening of the cell wall due to increased deposition of chitin (Vangalis *et al*., 2020), nuclear degradation and cell shrinkage, therefore resembling the incompatibility reactions described in other fungi (Dementhon *et al*., 2003; Glass & Dementhon, 2006; Gonçalves *et al*., 2017). Such reactions are mediated by allelic interactions between *sec-9*/*plp-1* (Heller *et al*., 2018) and *rcd-1* (Daskalov *et al*., 2019) in *N. crassa*, as well as different repertoires of multiple *het* loci among fungi (Saupe, 2000; Glass & Dementhon, 2006; Paoletti, 2016). However, a fundamental difference in *V. dahliae* is that a significant fraction of “incompatible” conidial fusions escapes allorecognition control under starvation, ensuring the formation of viable heterokaryons, in agreement with similar observations in the other asexual fungi *F. oxysporum* (Shahi *et al*., 2016) and *C. lindemuthianum* (Ishikawa *et al*., 2012).

Overall, our data favor the hypothesis of a trade-off between two mutually exclusive processes that both provide advantages to the adaptive evolution of asexual fungi, such as *V. dahliae*. On the one hand, genetic intermixing between individuals is subject to allorecognition control, which triggers cell death in the majority of “incompatible” fusions, possibly to protect these syncytial organisms from the spread of infectious or parasitic elements (Paoletti, 2016; Daskalov *et al*., 2017). On the other hand, rarity of nutrients induces cell fusion (Vangalis *et al*., 2021), and a fraction of the resulting heterokaryons overcome the imperfect incompatibility barriers hypothetically to grant populations with access to parasexuality and, therefore, genetic diversification that can be advantageous under stressful conditions. Even low levels of parasexual genomic reshuffling could have a marked effect on the genetic structure of fungal populations (Hoekstra, 1994) and could explain the signs of recombination observed in *V. dahliae* populations (Atallah *et al*., 2010; Milgroom *et al*., 2014), as well as the imperfect clonality of its VCGs (Papaioannou *et al*., 2013a). Parasexuality could also be critical for interspecific hybridization (Steensels *et al*., 2021) and horizontal gene transfer (Fitzpatrick, 2012; Soanes & Richards, 2014), which may have significant evolutionary roles. This is supported by our evidence of interspecific interactions between *V. dahliae* and *V. nonalfalfae*, similarly to those recently reported between *Colletotrichum* species (Mehta & Baghela, 2021), and the amphidiploid nature of the related hybrid species *V. longisporum*, presumed to have arisen parasexually between *V. dahliae* and other species (Karapapa *et al*., 1997; Inderbitzin *et al*., 2011).

Our study provides evidence that the molecular machinery of autophagy might be involved in the proposed trade-off as a link between starvation and the cell fate decision in *V. dahliae* heterokaryons. We found that Atg1- and Atg8-mediated autophagy participates in the incompatibility reaction, but not as a mediator of cell death. Instead, autophagy seems to act against destruction of “incompatible” heterokaryons, which is in line with previous studies of *P. anserina* that revealed accelerated cell death in autophagy-deficient mutants (Pinan-Lucarré *et al*., 2005). The autophagic machinery could function to protect “incompatible” fusions, e.g. by degrading incompatibility effectors, and survival of each heterokaryon may depend on the success of this process. Since cell fusion is coupled with starvation, which also induces autophagy (Feng *et al*., 2014), the majority of fused cells must have high levels of autophagic activity and this could function in favor of their survival.

Previous studies of the parasexual cycle (Pontecorvo & Roper, 1952; Hastie, 1964) and the hypotheses of parasexual horizontal gene transfer (Fitzpatrick, 2012; Soanes & Richards, 2014) predict exchange of genetic material between distinct nuclei within heterokaryons. Consistently, we observed frequent nuclear migration through the anastomosis bridge both in “compatible” and “incompatible” heterokaryons, commonly generating hyphae with genetically different nuclei. The resulting dikaryons, however, were eventually subjected to a nuclear division-triggered selective degradation process targeting the host nucleus of the affected cellular compartment. This process involves Atg1- and Atg8-mediated macroautophagy of the whole nucleus (i.e. macronucleophagy), which acts as a homeostatic mechanism ensuring a uninucleate status, both in hyphae and in fused cells of *V. dahliae*. This mechanism appears conserved between uninucleate fungi, as suggested by similar observations in *F. oxysporum* (Corral-Ramos *et al*., 2015). Despite the eventual degradation of the resident nucleus, however, the preceding phase of nuclear co-existence in the heterokaryon appears to provide a sufficient window of opportunity for genetic exchanges. These lead to mosaic “compatible” or “incompatible” mycelia consisting of sectors with either parental-type nuclei or novel ones resulting from direct nuclear fusion (i.e. karyogamy) or transfer of chromatin. The latter could involve transfer of chromosomes or chromosomal fragments by nuclear membrane-bound micronuclei, such as those observed in *C. lindemuthianum* (Roca *et al*., 2003) and *F. oxysporum* (Shahi *et al*., 2016).

This report contributes to a better understanding of the evolution of asexual fungi. Our findings have important implications for the interpretation of the traditional VCG data, which are commonly discussed under the assumption of genetic isolation between VCGs in asexual species. We propose the evolution of imperfect incompatibility systems in *V. dahliae* to preserve the ability of the species to engage in parasexuality. This conclusion suggests important roles of heterokaryosis and parasexuality in fungal evolution, and it welcomes future research aiming at a better understanding of the extent and modes of parasexual recombination achieved through this process. We propose *V. dahliae* as a particularly suitable organism for such analyses, and we expect future experimentation to be facilitated by the methodological toolkit for CRISPR/Cas9-mediated gene targeting that we optimized in this study to overcome experimental limitations in this and many other fungi.

## Acknowledgements

Plasmids pFC332 and pFC334 (U.H. Mortensen, Technical University of Denmark) were obtained from Addgene. This research is co-financed by Greece and the European Union (European Social Fund-ESF) through the Operational Programme «Human Resources Development, Education and Lifelong Learning» in the context of the project “Strengthening Human Resources Research Potential via Doctorate Research” (MIS-5000432), implemented by the State Scholarships Foundation (IKY).

## Author contributions

Conceptualization: I.A.P. and M.A.T.; Methodology: I.A.P., M.A.T., M.K. and V.V.; Investigation: V.V., I.A.P. and I.L.; Formal analysis: V.V. and I.A.P.; Supervision: I.A.P. and M.A.T.; Visualization: I.A.P. and V.V.; Writing – Original Draft: I.A.P. and V.V.; Writing – Review & Editing: I.A.P., M.A.T., M.K. and V.V.; Funding acquisition: M.K. and M.A.T.

## SUPPORTING INFORMATION

**Fig. S1.**
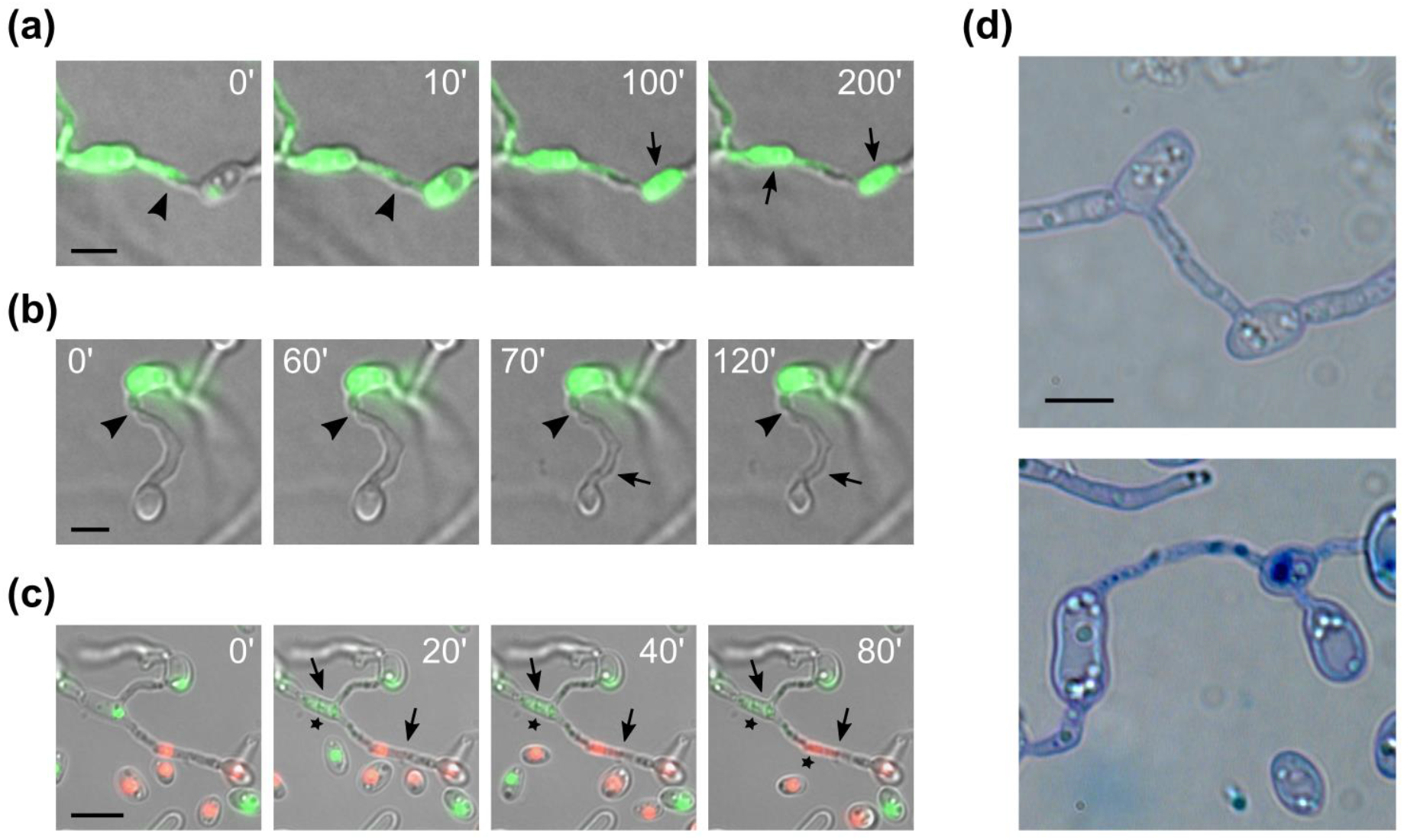
Typical features of incompatibility-triggered cell death in *V. dahliae*. (a) Induction of an incompatibility reaction after cytoplasmic mixing. Strains: PH sGFP and BB H1-sGFP. (b) Induction of an incompatibility reaction without prior cytoplasmic mixing. Strains: PH sGFP and BB. (c) Incompatibility-triggered cell death is characterized by nuclear degradation and cell shrinkage. Strains: PH H1-sGFP and Ls.17 H1-mCherry. In (a-c) arrowheads indicate the contact point of germlings, and arrows indicate cell shrinkage. (d) Staining of CAT-mediated fused cells using methylene blue. Viable fusions remain unstained (top), whereas post-fusion cell death permits the accumulation of the dye in the cytoplasm (bottom). Bars = 5 *μ*m.

**Fig. S2.**
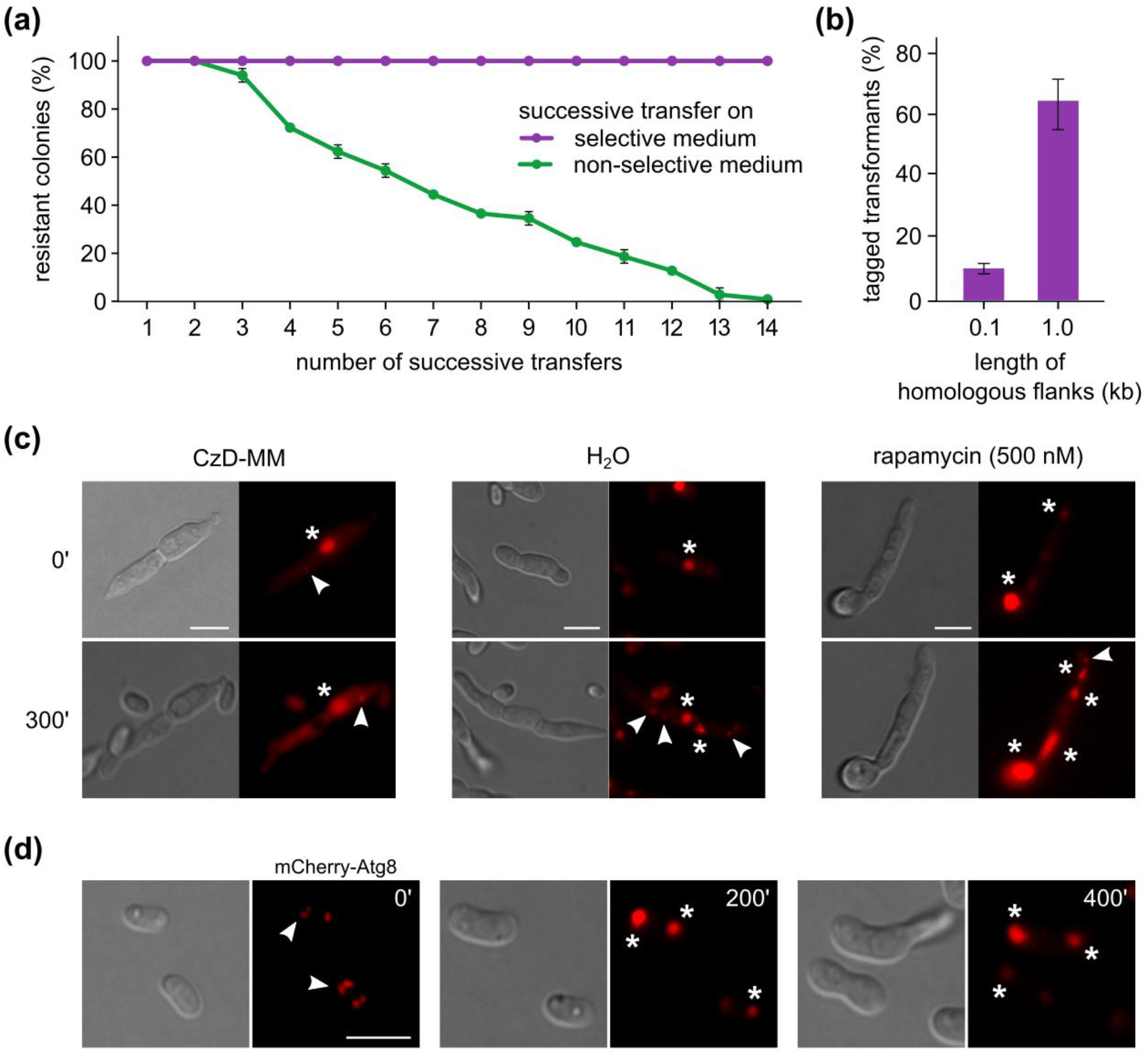
Fluorescent tagging of *V. dahliae* Atg8 by a CRISPR/Cas9-based system. (a) Stability of AMA1-containing plasmids under selective and non-selective conditions. Twenty-five randomly selected transformants of *V. dahliae* Ls.17 were single-spore purified and re-cultured every three days on selective (PDA supplemented with hygromycin B) and non-selective (PDA) medium. After each transfer, the percentage of resistant colonies to hygromycin B was determined. The experiment was performed in triplicate. Bars = SD. (b) Tagging efficiency of Atg8 with mCherry (strain Ls.17), using repair substrates with different lengths of homologous arms. The use of 0.1 kb-long arms yielded mostly chimeric fluorescent colonies consisting of tagged and non-tagged cell populations (13% properly tagged cells on average), in contrast to 1.0 kb-long arms that resulted in more homogeneous colonies (i.e. at least 90% of cells were properly tagged). The experiment was performed in triplicate for each repair construct, and at least 20 independent transformants were checked per replicate (using PCR and microscopy). Bars = SD. (c) Effect of autophagy-inducing conditions on the mCherry-Atg8 localization. Both starvation (i.e. incubation in water) and treatment with rapamycin (i.e. a TOR kinase inhibitor that induces autophagy) lead to the accumulation of the protein in autophagosomes and vacuoles, which indicates the induction of autophagy. (d) Subcellular localization of mCherry-Atg8 during germination in minimal medium. Prior to germination, the protein forms small globular foci (presumably autophagosomes, arrowheads), which seem to accumulate during germination in larger structures (vacuoles, asterisks), which is consistent with the expected participation of autophagy in conidial germination.

**Fig. S3.**
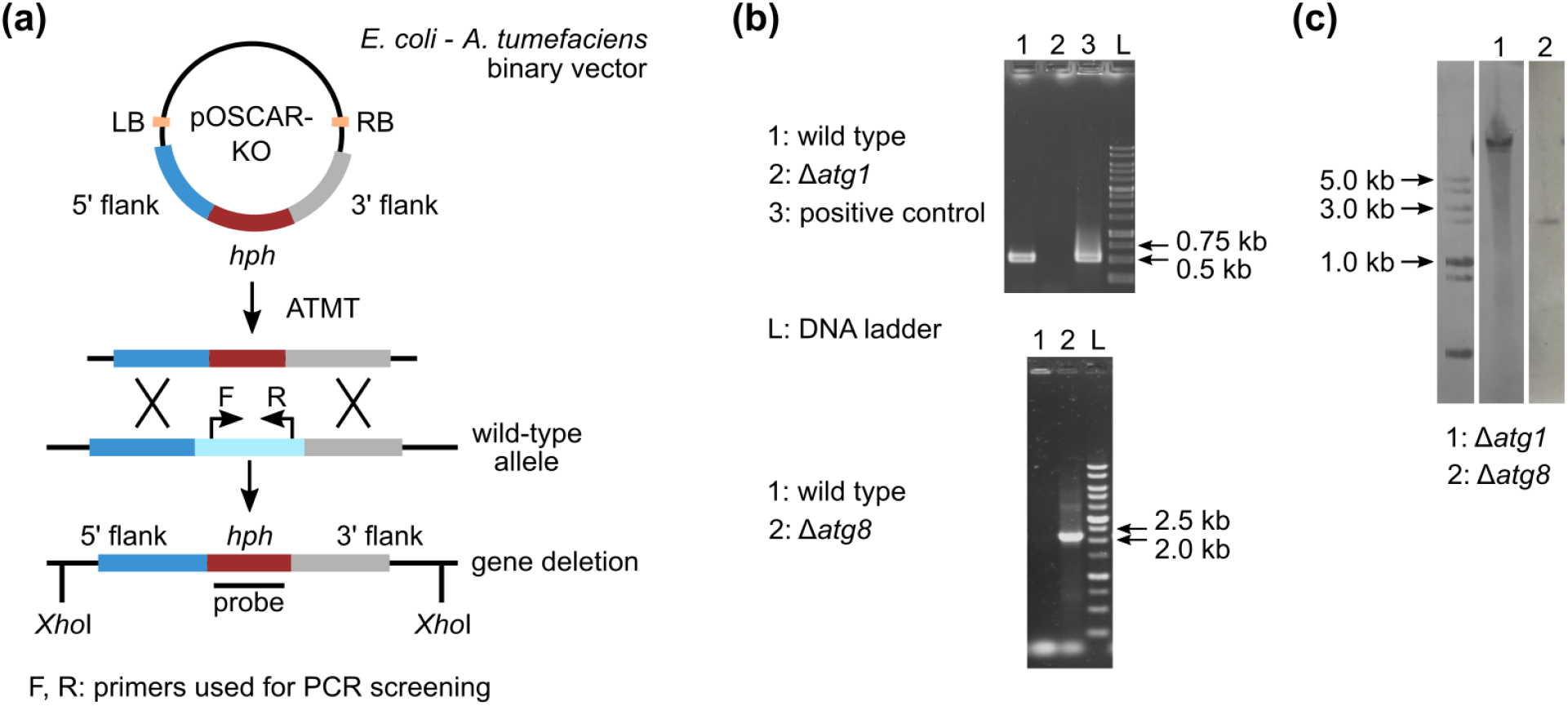
Construction and validation of *V. dahliae atg1* and *atg8* knockout mutants. (a) Schematic representation of the double homologous recombination-based strategy using *Agrobacterium tumefaciens*-mediated transformation for deletion of *atg1*. (b) Mutant validation by PCR using gene-specific (internal) primers (i.e. Vdatg1F/R) for the deletion of *atg1* and the combination of a gene-specific primer with a primer binding to the selection marker (i.e. atg8-gen-F/3flatg8R) for the CRISPR/Cas9-mediated disruption of *atg8*. (c) Validation of mutants as single integration events by Southern hybridization with the corresponding labeled selection marker cassettes as probes (i.e. *hph* for *atg1*, *neo*^R^ for *atg8*; genomic DNA of all strains was digested by *Xho*I).

**Fig. S4.**
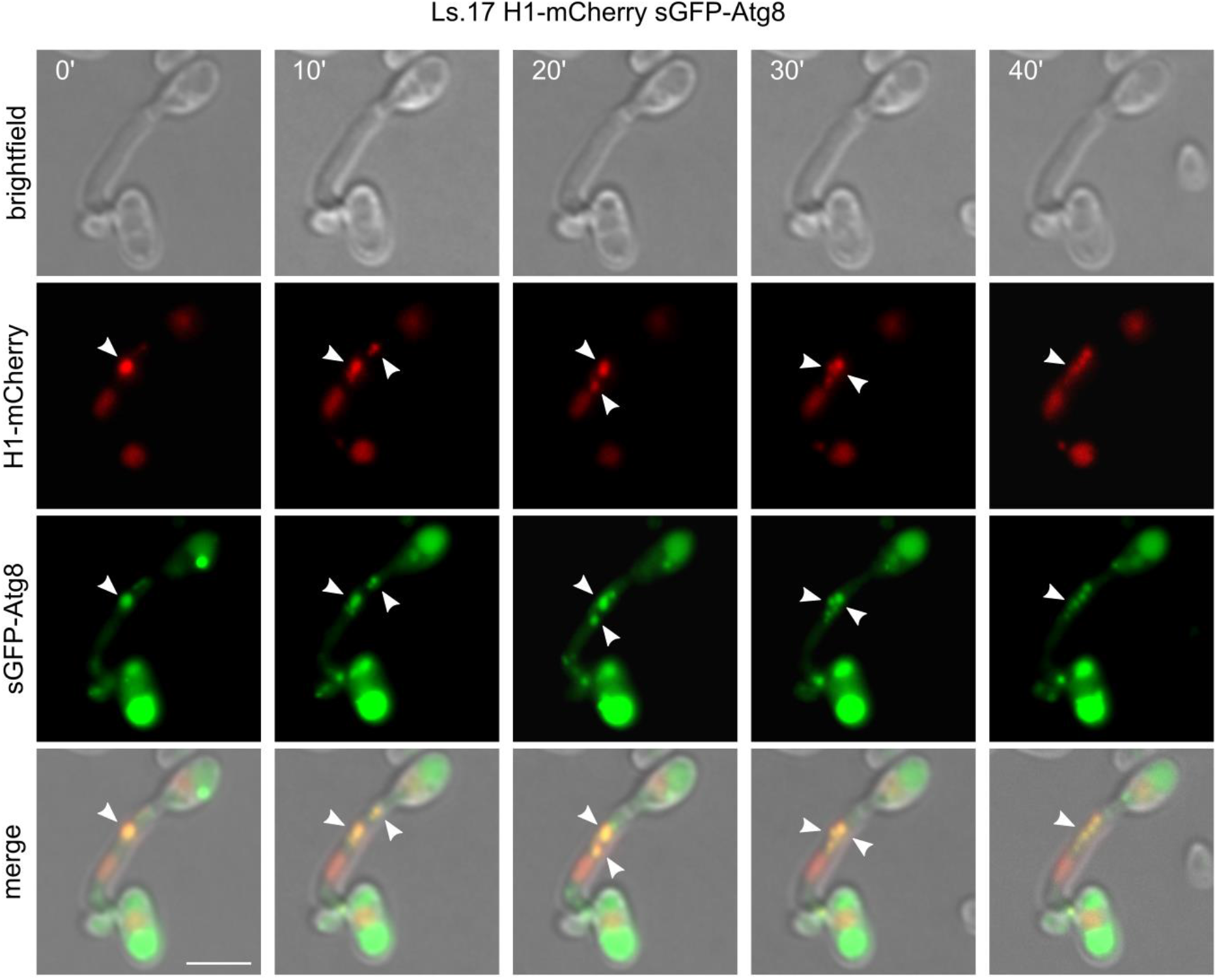
Atg8-mediated nuclear degradation in the strain Ls.17 H1-mCherry sGFP-Atg8. Atg8 is co-localized with the degrading nucleus during this process (arrowheads). Bars = 5 *μ*m.

**Fig. S5.**
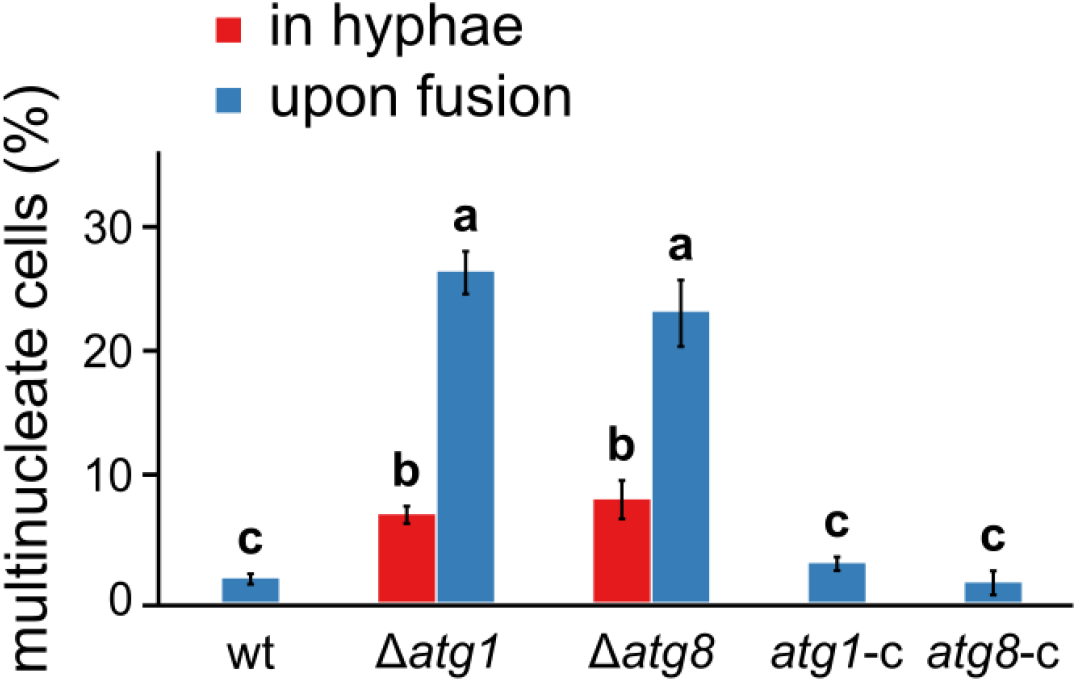
Frequencies of non-apical hyphal compartments and fused cells with more than one nuclei in complemented Δ*atg1* and Δ*atg8* knockout strains (by re-introducing the corresponding wild-type genes) in strain *V. dahliae* Ls.17. Each strain was tested in triplicate (n = 300 hyphal cells or 150 anastomoses per replicate). Statistical significance of differences was tested with one-way ANOVA followed by Tukey’s post-hoc test; bars with the same letter do not differ significantly (p-value > 0.05).

**Table S1.**
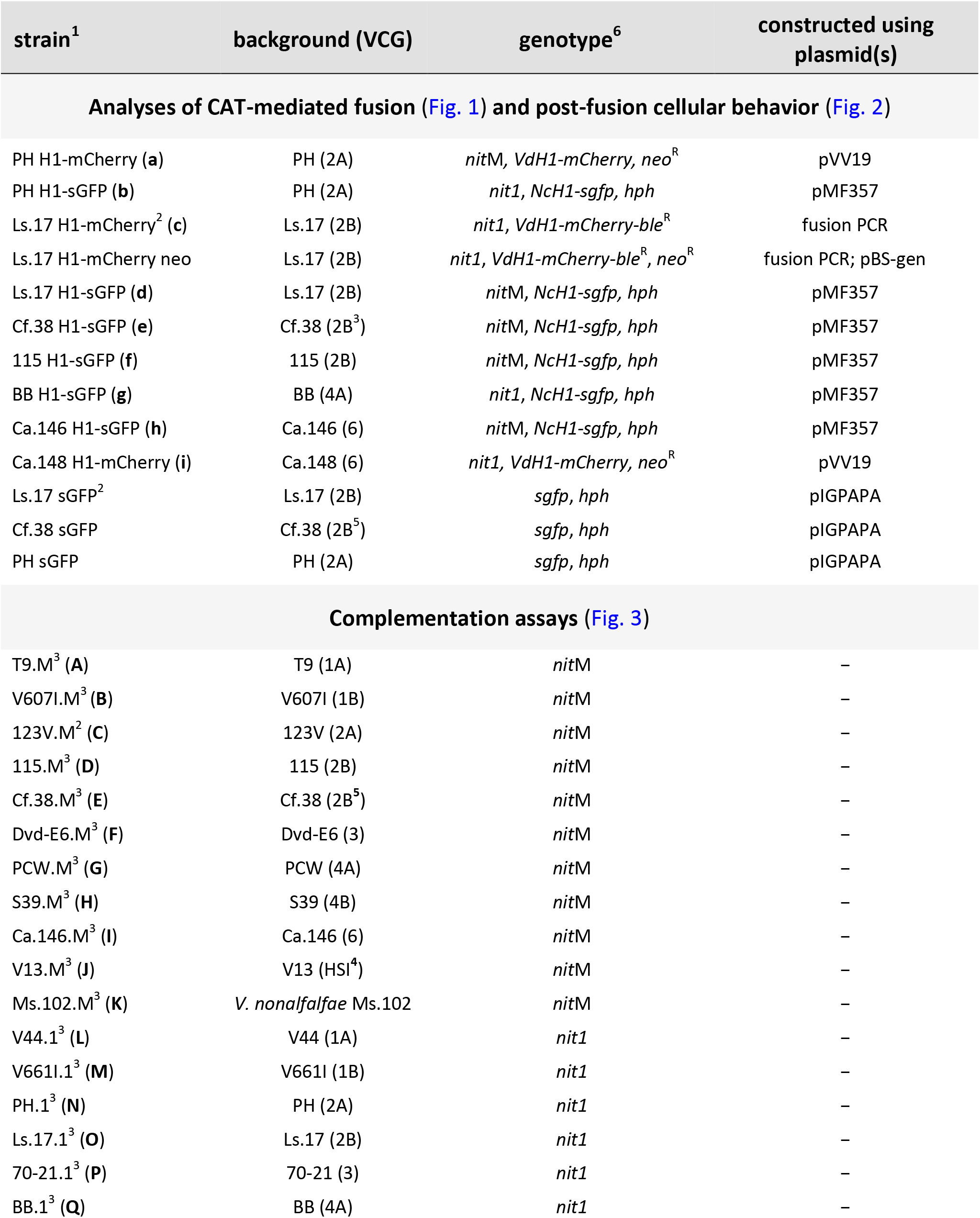
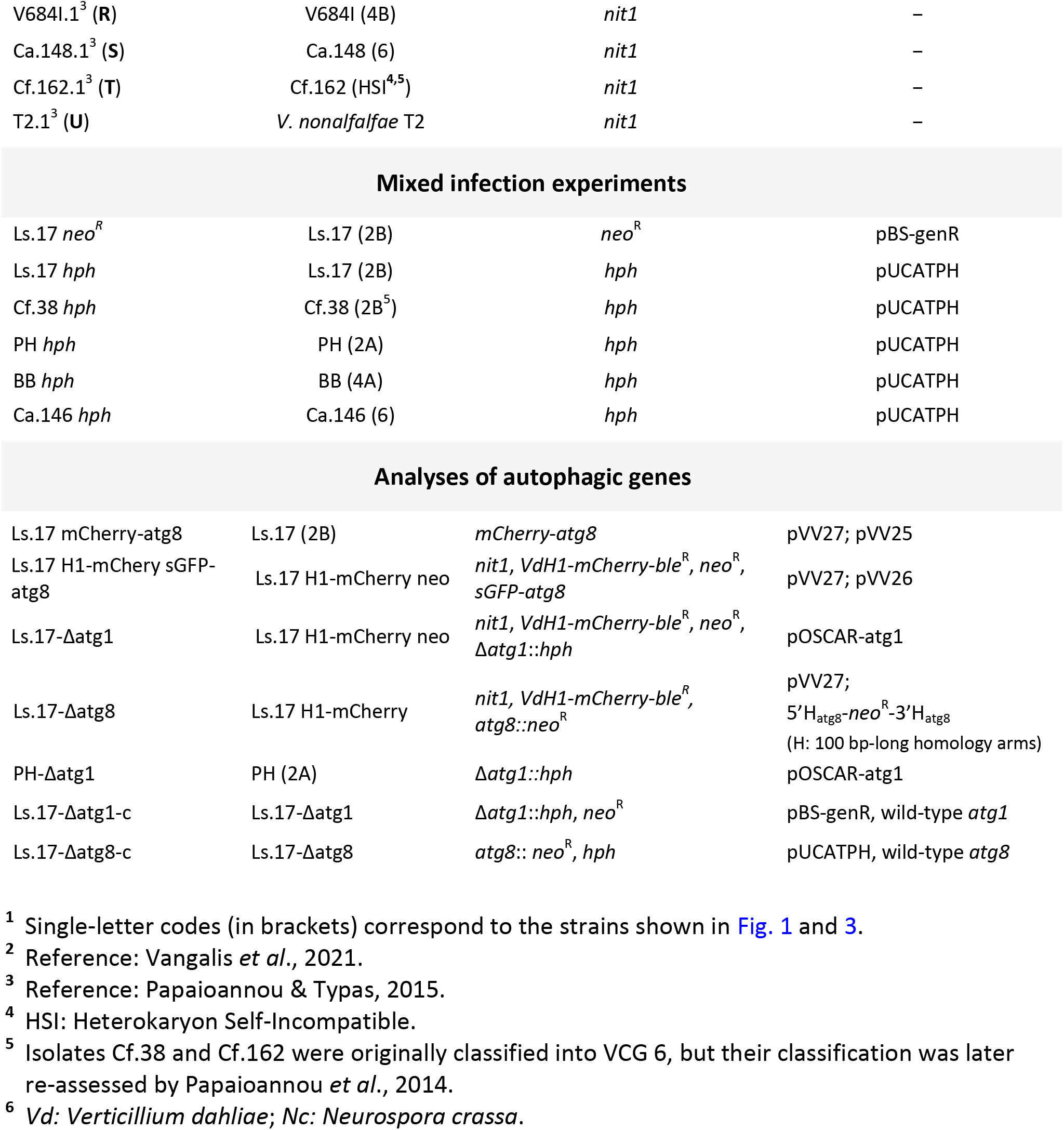
Verticillium dahliae strains used in this study.

| strain<sup>1</sup> | background (VCG) | genotype<sup>6</sup> | constructed using plasmid(s) |
| --- | --- | --- | --- |
| **Analyses of CAT-mediated fusion (Fig. 1) and post-fusion cellular behavior (Fig. 2)** |  |  |  |
| PH H1-mCherry (a) | PH (2A) | *nitM*, *VdH1-mCherry*, *neo*<sup>R</sup> | pVV19 |
| PH H1-sGFP (b) | PH (2A) | *nit1*, *NcH1-sgfp*, *hph* | pMF357 |
| Ls.17 H1-mCherry<sup>2</sup> (c) | Ls.17 (2B) | *nit1*, *VdH1-mCherry-ble*<sup>R</sup> | fusion PCR |
| Ls.17 H1-mCherry neo | Ls.17 (2B) | *nit1*, *VdH1-mCherry-ble*<sup>R</sup>, *neo*<sup>R</sup> | fusion PCR; pBS-gen |
| Ls.17 H1-sGFP (d) | Ls.17 (2B) | *nitM*, *NcH1-sgfp*, *hph* | pMF357 |
| Cf.38 H1-sGFP (e) | Cf.38 (2B<sup>3</sup>) | *nitM*, *NcH1-sgfp*, *hph* | pMF357 |
| 115 H1-sGFP (f) | 115 (2B) | *nitM*, *NcH1-sgfp*, *hph* | pMF357 |
| BB H1-sGFP (g) | BB (4A) | *nit1*, *NcH1-sgfp*, *hph* | pMF357 |
| Ca.146 H1-sGFP (h) | Ca.146 (6) | *nitM*, *NcH1-sgfp*, *hph* | pMF357 |
| Ca.148 H1-mCherry (i) | Ca.148 (6) | *nit1*, *VdH1-mCherry*, *neo*<sup>R</sup> | pVV19 |
| Ls.17 sGFP<sup>2</sup> | Ls.17 (2B) | *sgfp*, *hph* | pIGPAPA |
| Cf.38 sGFP | Cf.38 (2B<sup>5</sup>) | *sgfp*, *hph* | pIGPAPA |
| PH sGFP | PH (2A) | *sgfp*, *hph* | pIGPAPA |
| **Complementation assays (Fig. 3)** |  |  |  |
| T9.M<sup>3</sup> (A) | T9 (1A) | *nitM* | – |
| V607I.M<sup>3</sup> (B) | V607I (1B) | *nitM* | – |
| 123V.M<sup>2</sup> (C) | 123V (2A) | *nitM* | – |
| 115.M<sup>3</sup> (D) | 115 (2B) | *nitM* | – |
| Cf.38.M<sup>3</sup> (E) | Cf.38 (2B<sup>5</sup>) | *nitM* | – |
| Dvd-E6.M<sup>3</sup> (F) | Dvd-E6 (3) | *nitM* | – |
| PCW.M<sup>3</sup> (G) | PCW (4A) | *nitM* | – |
| S39.M<sup>3</sup> (H) | S39 (4B) | *nitM* | – |
| Ca.146.M<sup>3</sup> (I) | Ca.146 (6) | *nitM* | – |
| V13.M<sup>3</sup> (J) | V13 (HSI<sup>4</sup>) | *nitM* | – |
| Ms.102.M<sup>3</sup> (K) | *V. nonalfalfae* Ms.102 | *nitM* | – |
| V44.1<sup>3</sup> (L) | V44 (1A) | *nit1* | – |
| V661I.1<sup>3</sup> (M) | V661I (1B) | *nit1* | – |
| PH.1<sup>3</sup> (N) | PH (2A) | *nit1* | – |
| Ls.17.1<sup>3</sup> (O) | Ls.17 (2B) | *nit1* | – |
| 70-21.1<sup>3</sup> (P) | 70-21 (3) | *nit1* | – |
| BB.1<sup>3</sup> (Q) | BB (4A) | *nit1* | – |
| V684I.1<sup>3</sup> (R) | V684I (4B) | *nit1* | – |
| Ca.148.1<sup>3</sup> (S) | Ca.148 (6) | *nit1* | – |
| Cf.162.1<sup>3</sup> (T) | Cf.162 (HSI<sup>4,5</sup>) | *nit1* | – |
| T2.1<sup>3</sup> (U) | *V. nonalfalfae* T2 | *nit1* | – |
| Mixed infection experiments |  |  |  |
| Ls.17 *neo*<sup>R</sup> | Ls.17 (2B) | *neo*<sup>R</sup> | pBS-genR |
| Ls.17 *hph* | Ls.17 (2B) | *hph* | pUCATPH |
| Cf.38 *hph* | Cf.38 (2B<sup>5</sup>) | *hph* | pUCATPH |
| PH *hph* | PH (2A) | *hph* | pUCATPH |
| BB *hph* | BB (4A) | *hph* | pUCATPH |
| Ca.146 *hph* | Ca.146 (6) | *hph* | pUCATPH |
| Analyses of autophagic genes |  |  |  |
| Ls.17 mCherry-atg8 | Ls.17 (2B) | *mCherry-atg8* | pVV27; pVV25 |
| Ls.17 H1-mCherry sGFP-atg8 | Ls.17 H1-mCherry neo | *nit1*, *VdH1-mCherry-ble*<sup>R</sup>, *neo*<sup>R</sup>, *sGFP-atg8* | pVV27; pVV26 |
| Ls.17-Δatg1 | Ls.17 H1-mCherry neo | *nit1*, *VdH1-mCherry-ble*<sup>R</sup>, *neo*<sup>R</sup>, *Δatg1::hph* | pOSCAR-atg1 |
| Ls.17-Δatg8 | Ls.17 H1-mCherry | *nit1*, *VdH1-mCherry-ble*<sup>R</sup>, *atg8::neo*<sup>R</sup> | pVV27;<br>5′H<sub>atg8</sub>-*neo*<sup>R</sup>-3′H<sub>atg8</sub><br>(H: 100 bp-long homology arms) |
| PH-Δatg1 | PH (2A) | *Δatg1::hph* | pOSCAR-atg1 |
| Ls.17-Δatg1-c | Ls.17-Δatg1 | *Δatg1::hph*, *neo*<sup>R</sup> | pBS-genR, wild-type *atg1* |
| Ls.17-Δatg8-c | Ls.17-Δatg8 | *atg8::neo*<sup>R</sup>, *hph* | pUCATPH, wild-type *atg8* |

<sup>1</sup> Single-letter codes (in brackets) correspond to the strains shown in Fig. 1 and 3.
<sup>2</sup> Reference: Vangalis *et al.*, 2021.
<sup>3</sup> Reference: Papaioannou & Typas, 2015.
<sup>4</sup> HSI: Heterokaryon Self-Incompatible.
<sup>5</sup> Isolates Cf.38 and Cf.162 were originally classified into VCG 6, but their classification was later re-assessed by Papaioannou *et al.*, 2014.
<sup>6</sup> *Vd*: *Verticillium dahliae*; *Nc*: *Neurospora crassa*.

**Table S2.** List of plasmids constructed and used in this study.

| plasmid | backbone | description | source |
| --- | --- | --- | --- |
| **recombinant vector construction** |  |  |  |
| pUCATPH | - | *hph* | Lu *et al.*, 1994 |
| pMaM330 | pFA6a | *mCherry-myeGFP-kanMX6* | Khmelniskii *et al.*, 2015 |
| pSD1 | pBluescript II | $P_{gpdA}, P_{trpC}, neo^R$ | Nguyen *et al.*, 2008 |
| pBS-genR | pBluescript II | $neo^R$ | Vangalis *et al.*, 2020 |
| pFC332 | - | AMA1, *hph*, $P_{tef1}$-*SpCas9-Ttef1* | Nodvig *et al.*, 2015 |
| pFC334 | - | AMA1, *argB*, $P_{gpdA}$-HH-sgRNA backbone-HDV -$T_{trpC}$ | Nodvig *et al.*, 2015 |
| pOSCAR | pPZP-RCS2 | *A. tumafaciens* binary vector | Paz *et al.*, 2011 |
| **fluorescent labeling** |  |  |  |
| pMF357 | - | $P_{ccg1}$-*Nch1-sgfp*, *hph* | Ishikawa *et al.*, 2012 |
| pVV19 | pOSCAR | $P_{VdH1}$-*VdH1-mCherryFP-T<sub>tef1</sub>*, $neo^R$ | This study |
| pIGPAPA | - | $P_{icl}$-*sgfp-T<sub>nosI</sub>*, *hph* | Horwitz *et al.*, 1999 |
| **CRISPR/Cas9 mediated *atg8*-targeting** |  |  |  |
| pVV25 | pOSCAR | 5'$H_{Vdatg8}$-*sgfp*-3'$H_{Vdatg8}$<br>(H: 1.0 kb-long homology arms) | This study |
| pVV26 | pOSCAR | 5'$H_{Vdatg8}$-*mCherryFP*-3'$H_{Vdatg8}$<br>(H: 1.0 kb-long homology arms) | This study |
| pVV27 | pFC332 | $P_{gpdA}$-HH-crRNA for *Vdatg8*-HDV-$T_{trpC}$ | This study |
| ***atg1* deletion** |  |  |  |
| pOSCAR-atg1 | pOSCAR | 5'$H_{Vdatg1}$-*hph*-3'$H_{Vdatg1}$<br>(H: 2.0 kb-long homology arms) | This study |

**Table S3.**
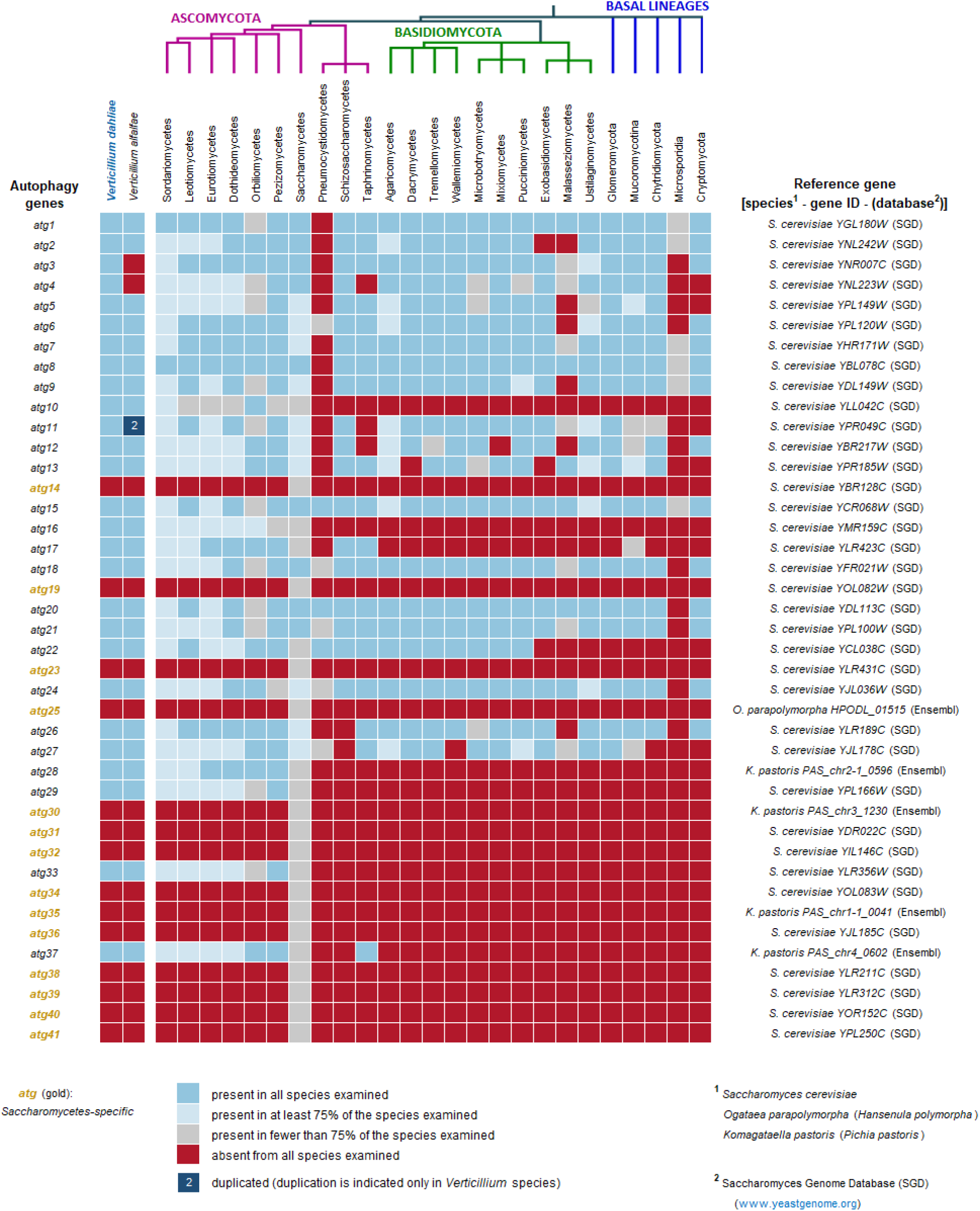
*Verticillium dahliae* homologs of autophagy genes.

**Table S4.**
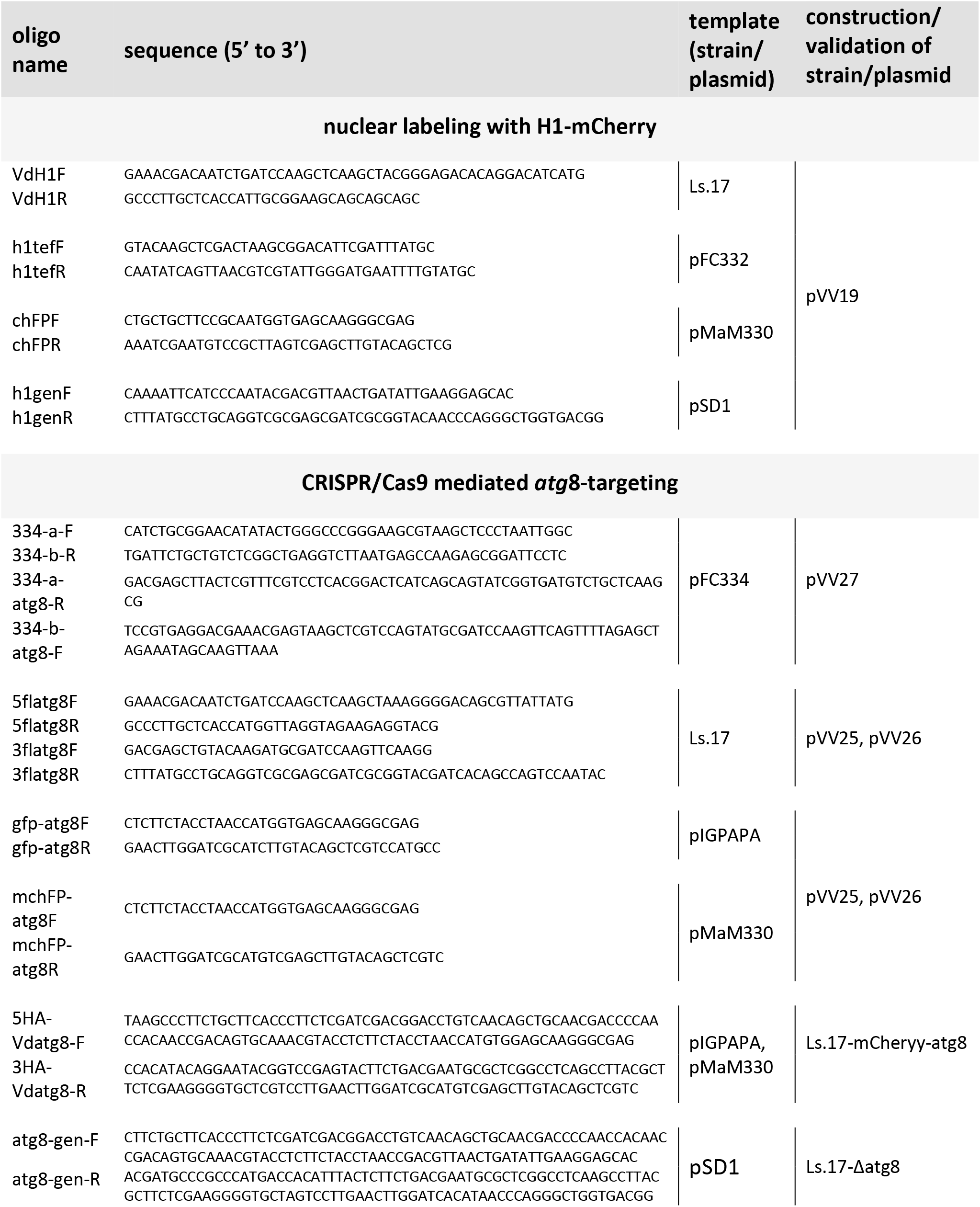
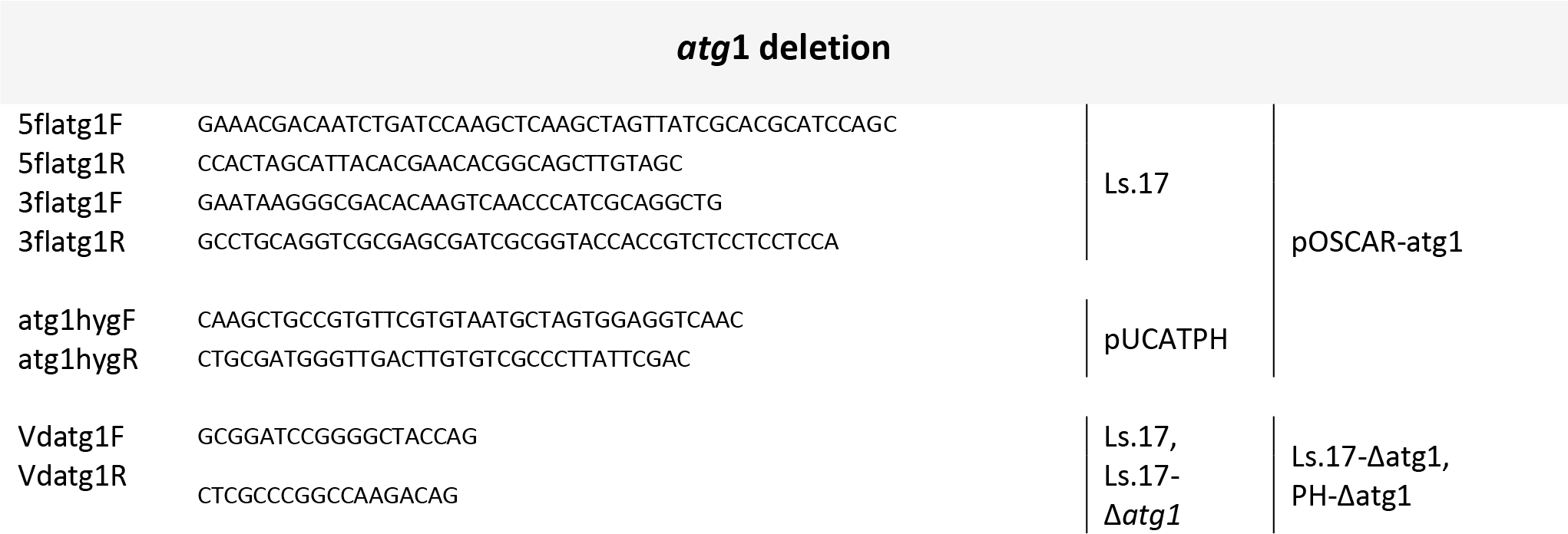
List of DNA oligonucleotides used in this study.

**Table S5.**
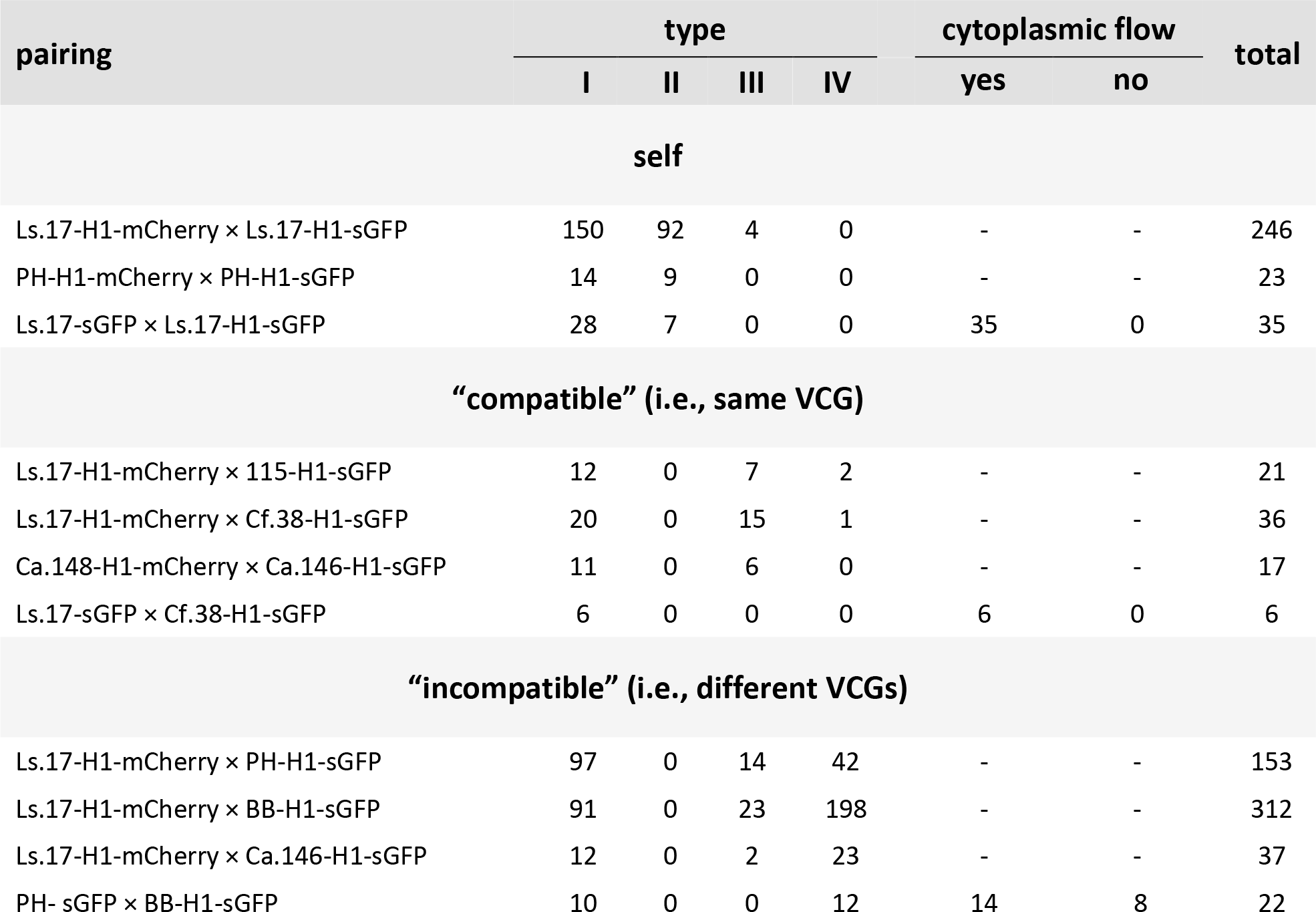
Cell fate and nuclear behavior (types I-IV, details in the text) of *V. dahliae* conidia/germlings involved in CAT-mediated fusion (summarized in Fig. 2).

**Table S6.** Results of the complementation tests summarized in Fig. 3. The vigor of each interaction is indicated by “+” (i.e. strong reaction characterized by dense prototrophic mycelial growth and extended production of microsclerotia), “+/-” (i.e. weak reaction with limited mycelial growth and/or restricted production of microsclerotia), or “-” (i.e. absence of prototrophic growth). Each pairing was performed in at least three replicates. Pairings that produced very limited prototrophic growth in only one of the replicates were regarded as negative (“-”).

| pairing | VCGs | assay |  |
| --- | --- | --- | --- |
|  |  | hyphal-based / VCG<br>(Fig. 3a,b) | CAT-based<br>(Fig. 3c,d) |
| T9.M × V44.1 | 1A × 1A | + | + |
| T9.M × V661I.1 | 1A × 1B | +/- | + |
| T9.M × PH.1 | 1A × 2A | - | +/- |
| T9.M × Ls.17.1 | 1A × 2B | - | - |
| T9.M × 70-21.1 | 1A × 3 | - | +/- |
| T9.M × BB.1 | 1A × 4A | - | +/- |
| T9.M × V684I.1 | 1A × 4B | - | +/- |
| T9.M × Ca.148.1 | 1A × 6 | - | +/- |
| T9.M × Cf.162.1 | 1A × HSI | - | - |
| T9.M × T2.1 | 1A × *V. nonalfalfae* | - | +/- |
| V607I.M × V44.1 | 1B × 1A | - | +/- |
| V607I.M × V661I.1 | 1B × 1B | +/- | +/- |
| V607I.M × PH.1 | 1B × 2A | - | - |
| V607I.M × Ls.17.1 | 1B × 2B | - | - |
| V607I.M × 70-21.1 | 1B × 3 | - | +/- |
| V607I.M × BB.1 | 1B × 4A | - | +/- |
| V607I.M × V684I.1 | 1B × 4B | - | - |
| V607I.M × Ca.148.1 | 1B × 6 | - | +/- |
| V607I.M × Cf.162.1 | 1B × HSI | - | +/- |
| V607I.M × T2.1 | 1B × *V. nonalfalfae* | - | - |
| 123V.M × V44.1 | 2A × 1A | - | - |
| 123V.M × V661I.1 | 2A × 1B | - | +/- |
| 123V.M × PH.1 | 2A × 2A | + | + |
| 123V.M × Ls.17.1 | 2A × 2B | - | + |
| 123V.M × 70-21.1 | 2A × 3 | - | - |
| 123V.M × BB.1 | 2A × 4A | - | +/- |
| 123V.M × V684I.1 | 2A × 4B | - | - |
| 123V.M × Ca.148.1 | 2A × 6 | - | +/- |
| 123V.M × Cf.162.1 | 2A × HSI | - | - |
| 123V.M × T2.1 | 2A × *V. nonalfalae* | - | +/- |
| 115.M × V44.1 | 2B × 1A | - | - |
| 115.M × V661I.1 | 2B × 1B | - | +/- |
| 115.M × PH.1 | 2B × 2A | - | +/- |
| 115.M × Ls.17.1 | 2B × 2B | +/- | + |
| 115.M × 70-21.1 | 2B × 3 | - | +/- |
| 115.M × BB.1 | 2B × 4A | - | +/- |
| 115.M × V684I.1 | 2B × 4B | - | +/- |
| 115.M × Ca.148.1 | 2B × 6 | - | +/- |
| 115.M × Cf.162.1 | 2B × HSI | - | +/- |
| 115.M × T2.1 | 2B × *V. nonalfalae* | - | - |
| Cf.38.M × V44.1 | 2B × 1A | - | - |
| Cf.38.M × V661I.1 | 2B × 1B | - | - |
| Cf.38.M × PH.1 | 2B × 2A | - | +/- |
| Cf.38.M × Ls.17.1 | 2B × 2B | + | + |
| Cf.38.M × 70-21.1 | 2B × 3 | - | +/- |
| Cf.38.M × BB.1 | 2B × 4A | - | +/- |
| Cf.38.M × V684I.1 | 2B × 4B | - | +/- |
| Cf.38.M × Ca.148.1 | 2B × 6 | - | +/- |
| Cf.38.M × Cf.162.1 | 2B × HSI | - | - |
| Cf.38.M × T2.1 | 2B × *V. nonalfalae* | - | - |
| Dvd-E6.M × V44.1 | 3 × 1A | - | - |
| Dvd-E6.M × V661I.1 | 3 × 1B | - | +/- |
| Dvd-E6.M × PH.1 | 3 × 2A | - | - |
| Dvd-E6.M × Ls.17.1 | 3 × 2B | - | +/- |
| Dvd-E6.M × 70-21.1 | 3 × 3 | + | +/- |
| Dvd-E6.M × BB.1 | 3 × 4A | - | +/- |
| Dvd-E6.M × V684I.1 | 3 × 4B | +/- | +/- |
| Dvd-E6.M × Ca.148.1 | 3 × 6 | - | - |
| Dvd-E6.M × Cf.162.1 | 3 × HSI | - | +/- |
| Dvd-E6.M × T2.1 | 3 × *V. nonalfalae* | - | - |
| PCW.M × V44.1 | 4A × 1A | - | +/- |
| PCW.M × V661I.1 | 4A × 1B | - | +/- |
| PCW.M × PH.1 | 4A × 2A | - | - |
| PCW.M × Ls.17.1 | 4A × 2B | - | - |
| PCW.M × 70-21.1 | 4A × 3 | - | + |
| PCW.M × BB.1 | 4A × 4A | + | +/- |
| PCW.M × V684I.1 | 4A × 4B | - | + |
| PCW.M × Ca.148.1 | 4A × 6 | - | +/- |
| PCW.M × Cf.162.1 | 4A × HSI | - | +/- |
| PCW.M × T2.1 | 4A × *V. nonalfalae* | - | +/- |
| S39.M × V44.1 | 4B × 1A | - | - |
| S39.M × V661I.1 | 4B × 1B | - | - |
| S39.M × PH.1 | 4B × 2A | - | - |
| S39.M × Ls.17.1 | 4B × 2B | - | +/- |
| S39.M × 70-21.1 | 4B × 3 | - | - |
| S39.M × BB.1 | 4B × 4A | - | - |
| S39.M × V684I.1 | 4B × 4B | +/- | +/- |
| S39.M × Ca.148.1 | 4B × 6 | - | +/- |
| S39.M × Cf.162.1 | 4B × HSI | - | - |
| S39.M × T2.1 | 4B × *V. nonalfalae* | - | - |
| Ca.146.M × V44.1 | 6 × 1A | - | +/- |
| Ca.146.M × V661I.1 | 6 × 1B | - | +/- |
| Ca.146.M × PH.1 | 6 × 2A | - | - |
| Ca.146.M × Ls.17.1 | 6 × 2B | - | - |
| Ca.146.M × 70-21.1 | 6 × 3 | - | +/- |
| Ca.146.M × BB. | 6 × 4A | - | - |
| Ca.146.M × V684I.1 | 6 × 4B | - | +/- |
| Ca.146.M × Ca.148.1 | 6 × 6 | + | + |
| Ca.146.M × Cf.162.1 | 6 × HSI | - | +/- |
| Ca.146.M × T2.1 | 6 × *V. nonalfalae* | - | +/- |
| V13.M × V44.1 | HSI × 1A | - | - |
| V13.M × V661I.1 | HSI × 1B | - | - |
| V13.M × PH.1 | HSI × 2A | - | - |
| V13.M × Ls.17.1 | HSI × 2B | - | - |
| V13.M × 70-21.1 | HSI × 3 | - | +/- |
| V13.M × BB.1 | HSI × 4A | - | +/- |
| V13.M × V684I.1 | HSI × 4B | - | - |
| V13.M × Ca.148.1 | HSI × 6 | - | - |
| V13.M × Cf.162.1 | HSI × HSI | - | - |
| V13.M × T2.1 | HSI × *V. nonalfalfae* | - | - |
| Ms.102.M × V44.1 | *V. nonalfalfae* × 1A | - | - |
| Ms.102.M × V661I.1 | *V. nonalfalfae* × 1B | - | +/- |
| Ms.102.M × PH.1 | *V. nonalfalfae* × 2A | - | - |
| Ms.102.M × Ls.17.1 | *V. nonalfalfae* × 2B | - | - |
| Ms.102.M × 70-21.1 | *V. nonalfalfae* × 3 | - | +/- |
| Ms.102.M × BB.1 | *V. nonalfalfae* × 4A | - | +/- |
| Ms.102.M × V684I.1 | *V. nonalfalfae* × 4B | - | +/- |
| Ms.102.M × Ca.148.1 | *V. nonalfalfae* × 6 | - | +/- |
| Ms.102.M × Cf.162.1 | *V. nonalfalfae* × HSI | - | - |
| Ms.102.M × T2.1 | *V. nonalfalfae* | +/- | +/- |

## Methods S1 Protocols for nuclear and cytoplasmic fluorescent labeling

Nuclei of *V. dahliae* strains were labeled with either sGFP- or mCherry-tagged histone H1. Carboxy-terminal tagging of *V. dahliae* histone H1 with the red fluorescent protein mCherry was performed using either a fusion PCR strategy that we have previously described (Vangalis *et al*., 2021) or a binary vector (pVV19) suitable for *Agrobacterium tumefaciens*-mediated transformation (ATMT).

The first method involved the generation of two PCR amplicons using the genomic DNA of *V. dahliae* isolate Ls.17 as template, containing the endogenous promoter and the coding region of the histone H1 gene (VDAG_09854), and its transcription termination sequence, respectively. The *mCherry* coding sequence and the phleomycin-resistance (*ble*^R^) cassette were amplified from plasmid pAN8.1-mCherry (Ruiz-Roldán *et al*., 2010). The two genomic amplicons and the *mCherry*-containing cassette were mixed in a PCR reaction without primers, which was followed by a second reaction that included the first PCR product as template and the primer pair VdH1FusF/VdH1FusR (Table S4). The fusion PCR product was then used to transform protoplasts of *V. dahliae* strains, according to our previously reported protocol (Vangalis *et al*., 2020).

Alternatively, we constructed plasmid pVV19 for mCherry-tagging of histone H1 by amplifying the following fragments with the Herculase II Fusion DNA polymerase (Agilent, Santa Clara, CA, USA) (all PCR primers are listed in Table S4): i. a 1,972 bp-long fragment from genomic DNA of *V. dahliae* strain Ls.17, containing the coding region of the histone H1 gene (VDAG_09854) excluding the stop codon, and a 1,000 bp-long upstream genomic fragment presumably including the endogenous promoter; ii. a 717 bp-long fragment from plasmid pMaM330 (Khmelinskii *et al*., 2016) containing the *mCherry* gene; iii. a 489 bp-long fragment from plasmid pFC332 (Nødvig *et al*., 2015) containing the transcription termination sequence of the *Aspergillus nidulans tef1* gene; and iv. a 1,648 bp-long fragment from plasmid pBS-genR (Vangalis *et al*., 2020) containing the G418-resistance (*neo*^R^) cassette. The four PCR amplicons were subjected to an *in vitro* DNA assembly reaction in the backbone of plasmid pOSCAR (Paz *et al*., 2011) using the NEBuilder HiFi DNA Assembly Master Mix kit (New England Biolabs, Ipswich, MA, USA), according to the manufacturer’s recommendations. The resulting vector was used for ATMT of *V. dahliae* strains, according to our recently reported protocol (Vangalis *et al*., 2020). Nuclear labeling of *V. dahliae* strains with heterologous sGFP-tagged histone H1 was performed using plasmid pMF357 (Ishikawa *et al*., 2012). This contains a fusion construct of the *sgfp* gene to the *Colletotrichum lindemuthianum* histone H1 gene and the *hph* cassette that confers resistance to hygromycin B. This plasmid was used for transformation of *V. dahliae* protoplasts (Vangalis *et al*., 2020).

*Verticillium dahliae* strains with cytoplasmic sGFP were constructed using plasmid pIGPAPA (Horwitz *et al*., 1999), which carries the *sgfp* gene under the control of the *Neurospora crassa ICL* promoter and the *hph* cassette, to transform protoplasts of the desired *V. dahliae* strains (Vangalis *et al*., 2020).

## Methods S2 Protocols for hyphal- and CAT-based complementation assays

In this study we used the traditional (VCG) complementation assay, which is hyphal-based, as well as a newly developed assay that involves CAT-mediated fusion of conidia/germlings of the interacting strains. For both methods, we used the same set of previously characterized nitrate non-utilizing *nit* mutants (*nit1* and *nit*M; with reversion rates lower than 1.0 × 10^-7^) of a collection of *V. dahliae* isolates of all VCGs (Table S1) (Papaioannou *et al*., 2014; Papaioannou & Typas, 2015; Vangalis *et al*., 2021).

In the VCG assay, sparse mycelial growth of complementary *nit* mutants on co-inoculated minimal medium (MM) plates enables mycelial confrontation of the paired isolates on the agar. When this interaction leads to heterokaryotic prototrophic growth along the confrontation zone (Fig. 3a), the corresponding strains are considered “compatible” and assigned to the same Vegetative Compatibility Group (VCG) (Joaquim & Rowe, 1990). We performed all VCG assays on Czapek-Dox MM agar (Vangalis *et al*., 2021) and scored plates for prototrophic growth four weeks after inoculation, according to standard procedures and previously described criteria (Papaioannou *et al*., 2013; Papaioannou & Typas, 2015). Each pairing was performed in triplicate and all used *nit* mutants were also inoculated separately on MM agar (in triplicate) to serve as controls.

We also developed and assessed in this study a CAT-based complementation assay of *nit* mutants. This method is based on pairing conidia of the desired *V. dahliae* strains under conditions that are favorable for CAT formation, and then transferring the samples to MM to select for heterokaryotic prototrophic growth (Fig. 3c). For this, we prepared a fresh conidial suspension (in water) from a 7-day-old PDA culture of each strain, and we diluted each suspension in CAT medium (0.75 g/l β-glycerophosphate disodium salt · 5H_2_O) to a final concentration of 1.0 × 10^6^ conidia/ml. The desired suspensions were then mixed (1:1) for each pairing, and 100 *μ*l of each mixture were transferred to a well of a polystyrene 96-well flat-bottom plate (2620/S from Kartell, Noviglio, Italy), with a cellophane disk placed at the bottom of the well. The plates were incubated at 24°C (in the dark) for 72 h before the cellophane disks were aseptically transferred to the surface of Czapek-Dox MM agar. The plates were scored for prototrophic growth on the cellophane disks over four weeks after inoculation, while they were incubated at 24°C (in the dark). Each pairing was performed in triplicate and all tested *nit* mutants were also subjected to the same procedure individually (in triplicate) as controls.

Presumed heterokaryons generated by both methods were randomly selected for further characterization to validate their heterokaryotic nature and exclude the possibilities of prototrophic growth because of marker reversion or cross-feeding of non-fused individuals (Typas & Heale, 1976). For this, they were sub-cultured on selective medium (MM), conidial suspensions were prepared from these cultures, and approx. 1.0 × 10^4^ conidia were spread on fresh MM. Conidia of *V. dahliae* are always unicellular and predominantly uninucleate (99.7% in the wild-type isolate Ls.17 according to our observations), while the rare bi- and multi-nucleate conidia are considered strictly homozygous due to their origin from the single nucleus of a phialide (Hastie, 1967; 1968). Therefore, a heterokaryon is expected to grow on selective medium due to genetic complementation between its genetically distinct nuclei, whereas its conidia would be of one of the parental types, and would thus remain unable to grow under selection for prototrophy.

**Movie S1** Establishment of cytoplasmic continuity following CAT-mediated fusion in an “incompatible” pairing (PH sGFP × BB H1-sGFP).

**Movie S2** Time-lapse imaging of a fusion event between the “incompatible” strains PH sGFP × BB H1-sGFP (cell wall staining: calcofluor white).

**Movie S3** Time-lapse imaging of a self-fusion of strain Ls.17 H1-mCherry sGFP-*atg8*. The host nucleus is surrounded by a ring-like Atg8-containing structure and eventually degraded.

**Movie S4** Time-lapse imaging of a self-fusion of strain Ls.17 H1-mCherry Δ*atg1*. Both nuclei of the dikaryon remain viable following nuclear migration.

**Movie S5** Nuclear division in a hyphal compartment of strain Ls.17 H1-mCherry Δ*atg8* results in the formation of a binucleate cell.

**Movie S6** Nuclear division in a non-apical hyphal compartment of strain Ls.17 H1-mCherry is followed by degradation of one of the daughter nuclei. Only one nucleus remains viable in the compartment.

## Notes

### Competing Interest Statement

The authors have declared no competing interest.

